# A gut meta-interactome map reveals modulation of human immunity by microbiome effectors

**DOI:** 10.1101/2023.09.25.559292

**Authors:** Veronika Young, Bushra Dohai, Thomas C. A. Hitch, Patrick Hyden, Benjamin Weller, Niels S. van Heusden, Deeya Saha, Jaime Fernandez Macgregor, Sibusiso B. Maseko, Chung-Wen Lin, Mégane Boujeant, Sébastien A. Choteau, Franziska Ober, Patrick Schwehn, Simin Rothballer, Melina Altmann, Stefan Altmann, Alexandra Strobel, Michael Rothballer, Marie Tofaute, Matthias Heinig, Thomas Clavel, Jean-Claude Twizere, Renaud Vincentelli, Marianne Boes, Daniel Krappmann, Claudia Falter, Thomas Rattei, Christine Brun, Andreas Zanzoni, Pascal Falter-Braun

## Abstract

The molecular mechanisms by which the gut microbiome influences human health remain largely unknown. Pseudomonadota is the third most abundant phylum in normal gut microbiomes. Several pathogens in this phylum can inject so-called virulence effector proteins into host cells. We report the identification of intact type 3 secretion systems (T3SS) in 5 - 20% of commensal Pseudomonadota in normal human gut microbiomes. To understand their functions, we experimentally generated a high-quality protein-protein meta-interactome map consisting of 1,263 interactions between 289 bacterial effectors and 430 human proteins. Effector targets are enriched for metabolic and immune functions and for genetic variation of microbiome-influenced traits including autoimmune diseases. We demonstrate that effectors modulate NF-κB signaling, cytokine secretion, and adhesion molecule expression. Finally, effectors are enriched in metagenomes of Crohn’s disease, but not ulcerative colitis patients pointing toward complex contributions to the etiology of inflammatory bowel diseases. Our results suggest that effector-host protein interactions are an important regulatory layer by which the microbiome impacts human health.

## MAIN

The host-associated microbiota influences human health in complex host genetics-dependent ways^1,2^. Especially intestinal microbes positively and negatively affect the risk for several complex diseases ranging from inflammatory bowel disease (IBD)^1^ and asthma^3^ to metabolic^4^ and neurodegenerative diseases^5^. Members of the bacterial phylum Pseudomonadota (previously: Proteobacteria^6^) are prevalent in the human gut microbiome and their occurrence is influenced by dietary ingredients such as fat and artificial sweeteners^7^. Unique features of this phylum are the type-3, type-4, and type-6 secretion systems (TxSS) that enable the injection of bacterial proteins directly into the host cytosol. The presence of T3SS has been classically associated with pathogen virulence^8^. In the plant kingdom, however, important mutualistic microbes also communicate with the host via effector proteins to establish cohabitation and elicit host-beneficial effects^9^. We therefore wondered if commensal Pseudomonadota in the healthy human gut microbiome possess host-directed secretion systems.

### T3SS are common in the normal human gut microbiome

Because of the higher quality and completeness of genome assemblies from cultured strains compared to metagenome-assembled genomes (MAGs), we first evaluated Pseudomonadota strains from gut and stool samples that were collected, among others, by the human microbiome project and were available from culture collections. Using EffectiveDB^10^, a widely used tool for secretion system identification, we detected complete T3SS in 44 of the 77 reference strain genomes (Extended Data Table 1). To expand the scope, we analyzed genomes of 4,752 distinct strains, representing all major phyla from the human gut that had been isolated by the human gastrointestinal bacteria genome collection (HBC)^11^, and the Unified Human Gastrointestinal Genome (UHGG) collection^12,13^. Of the 2,272 Gram-negative strains, 478 (21%) had complete T3SS (Fig. 1a); similar proportions have T4SS (527) and T6SS (719), both of which can also deliver effectors into host cells but also have other functions (Extended Data Fig. 1 and Extended Data Table 1)^14^. Together 729 of the 2,272 Gram-negative strains, *i.e.*, 34%, have at least one host-directed secretion system. Because culturing can bias the relative proportions of taxa, we sought to confirm the presence of T3SS in commensal microbiota using metagenome datasets. From 16,179 Pseudomonadota MAG bins with high or intermediate genome quality^15,16^, 770, i.e., 5%, encoded complete T3SS (Fig. 1a and Extended Data Table 1). Notably, we only identified T3SS in Gammaproteobacteria, whereas no secretion systems were found in the Beta- or Epsilonproteobacteria in the datasets, except for a few *Helicobacter* strains. It is unclear if gut commensal strains in these orders lack T3SS, or if the systems differ from those of the better-characterized Gammaproteobacteria and they were missed by the algorithm. Across the analyses, T3SS were identified in strains of multiple genera and were especially common among *Escherichia* (Fig. 1b and Extended Data Table 1). Notably, a recent *in vivo* profiling study of human digestive tracts using *in situ* sampling found *Escherichia* as the genus that was most significantly enriched in intestinal over stool samples^17^. Of the T3SS-positive (T3SS^+^) species, 24 matched representatives in two cohorts of a dataset provided by the Weizmann Institute of Science (WIS cohorts)^18^. 59.4% of individuals in the Israeli cohort and 47.1% in the Dutch cohort had potentially T3SS^+^ species in their gut microbiome, with relative abundances of 0.80% and 0.48%, respectively (Fig. 1c). The most common T3SS^+^ species in both cohorts was *Escherichia coli*, appearing within 54% and 45% of individuals, respectively. Overall, T3SS^+^ strains constitute a substantial proportion of commensal Pseudomonadota and are common in normal human gut microbiomes. We therefore aimed to understand the functions of T3SS-delivered effector proteins of commensal strains.

**Fig. 1.**
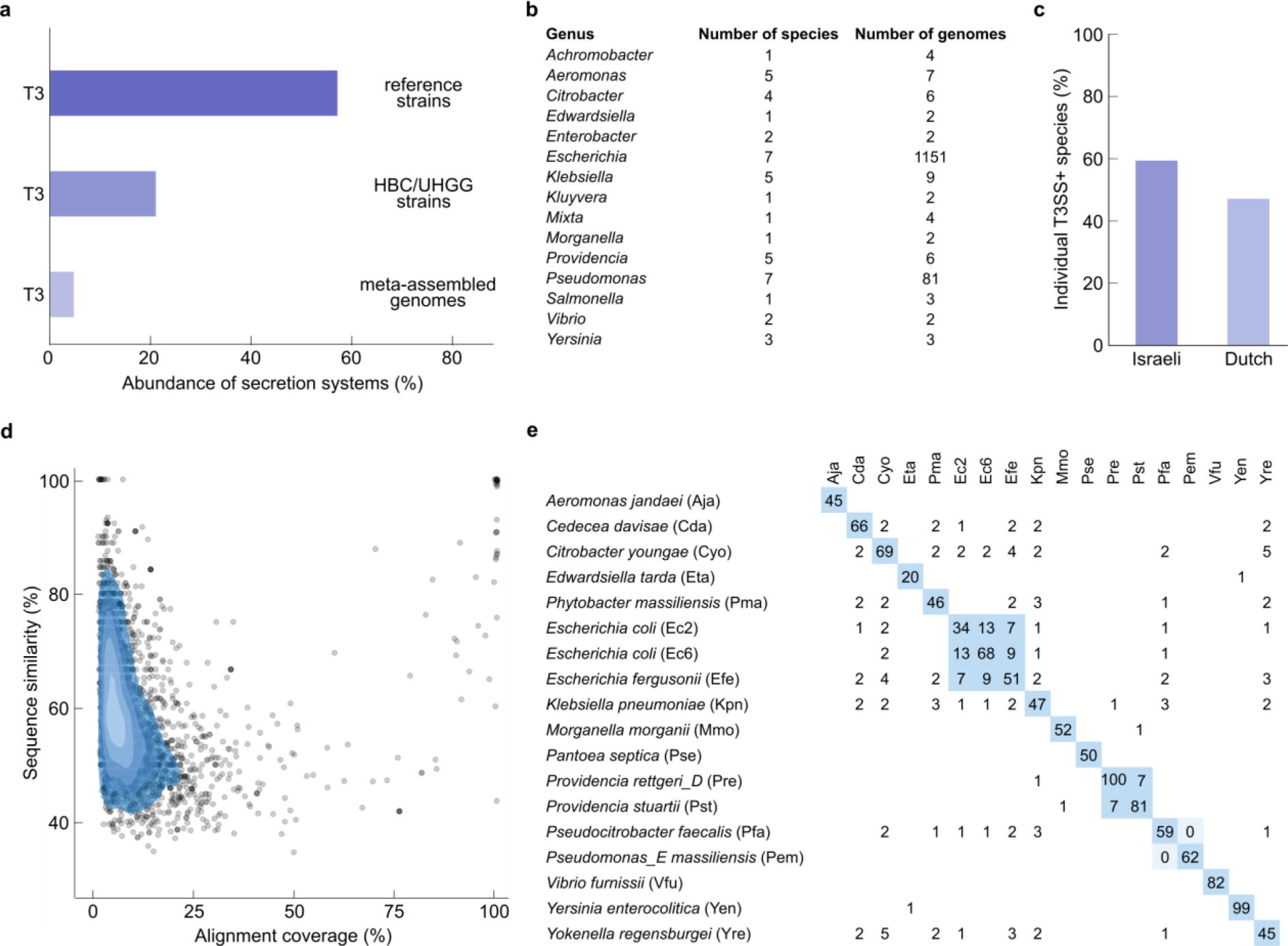
| T3SS in commensal bacterial species in the gut microbiome. **a**, Proportion of Pseudomonadota genomes encoding complete T3SS among 77 reference strains of human intestinal and stool samples, in a collection of 4,475 strains isolated from normal human guts, and in meta-assembled genomes (MAG) of normal human guts. **b**, Most abundant genera and identified number of species and genomes encoding complete T3SS from the samples in **a**. **c**, Proportion of individuals in two human cohorts containing T3SS encoding microbial species. **d**, Similarity of 3,002 candidate effector-substrates for T3SS identified from commensal reference strains with 1,195 effectors from pathogenic microbes across the range of alignment coverages. **e**, Selection of 18 commensal Pseudomonadota strains with dissimilar effector complements used for subsequent functional analyses. Numbers indicate the count of shared effectors at >90% mutual sequence similarity across 90% common sequence length among the indicated strains. Full data for all panels in Extended Data Table 1.

### Commensal effectors are unrelated to known pathogen effectors

To identify gut microbiome-encoded effectors we used a combination of three complementary machine learning models^19–21^ and considered 3,002 effector candidates from the 44 reference strains that were most confidently predicted by all tools (Extended Data Table 2). In addition, we identified 186 putative effectors in the 770 T3SS^+^ MAGs (Extended Data Table 2). As T3SS and substrate effectors are best known for their role in supporting a pathogenic lifestyle, we investigated if the commensal bacterial effectors share sequence similarity with 1,638 known T3SS effectors from pathogens^22^. Only 17 of 3,002 (0.5%) effectors from strains and 6 of 186 (3%) from MAGs, respectively, showed extended high sequence similarity (≥ 90% sequence similarity across ≥ 90% length) to known pathogen effectors; lowering the thresholds to 50% similarity across 75% length only marginally increased the numbers to 34 (1%) and 7 (4%), respectively (Fig. 1d and Extended Data Table 2). The largest number of commensal effectors with similarity to pathogenic effectors were found in the genomes of *Escherichia albertii* (12 effectors with 67% to 98% identity) and *Yersinia enterocolitica* (10 effectors at > 98% identity). The fact that all such pathogen-similar commensal effectors were found in different species, of which some even belong to a different order than the respective pathogen, suggests that non-pathogenic microbes participate in the horizontal gene transfer of effectors^23,24^. This is supported by the observation that only a few pathogen-similar effectors were found among the approximately 20 - 80 effectors of each strain. Of the six pathogen-similar effectors found in MAGs, all but one matched the identified family of the pathogen from which they were initially identified (Extended Data Fig. 2 and Extended Data Table 2). Plausibly, these effectors originate from pathogens, or their relatives that were likely present in some samples. Jointly, the data show that effector complements of commensal bacteria are distinct from those of pathogens, thereby suggesting functions outside of the pathogen lifestyle.

### A microbiome-host protein-protein meta-interactome map

To investigate the functions of commensal effectors, we cloned effector ORFs for experimental studies from 18 bacterial strains with diverse effector complements (Fig. 1e and Extended Data Fig. 1). We successfully PCR-cloned 786 ORFs for the 1,300 encoded effectors (60.2%) and 173 of 186 effector ORFs from MAG bins (meta-effectors) following chemical synthesis (Fig. 2a). Thus, 959 sequence-verified full-length effector ORFs were assembled as the human microbiome effector ORFeome (HuMEOme_v1) (Extended Data Table 2). With these, we conducted binary interactome (contactome) network mapping against the human ORFeome9.1 collection encoding 18,000 human gene products using a stringent multi-assay mapping pipeline^25^. In the main screen by yeast-2-hybrid (Y2H), we identified 1,071 interactions constituting the <u>hu</u>man-<u>m</u>icrobiome <u>m</u>eta-interactome main dataset (HuMMI_MAIN_) (Fig. 2b,c). To assess sampling sensitivity^26^, i.e., saturation of the screen, we conducted three additional repeats of 290 randomly picked effectors and 1,440 human proteins, which yielded 39 verifiable interactions constituting the HuMMI repeat subset (HuMMI_RPT_). The saturation curve indicates that the single main screen has a sampling sensitivity of ∼32% (Fig. 2d). Last, to address how effector sequence similarity affects their interaction profiles we conducted a homolog screen. Effectors were grouped if they shared ≥ 30% sequence identity (Extended Data Table 2) and all effectors of one group were experimentally tested against the union of their human interactors. The resulting dataset (HuMMI_HOM_) contains 398 verified interactions, of which 179 were not found in the other screens. Altogether, HuMMI contains 1,263 unique verified interactions between 289 effectors and 430 human proteins (Fig. 2b,c and Extended Data Table 3).

**Fig. 2.**
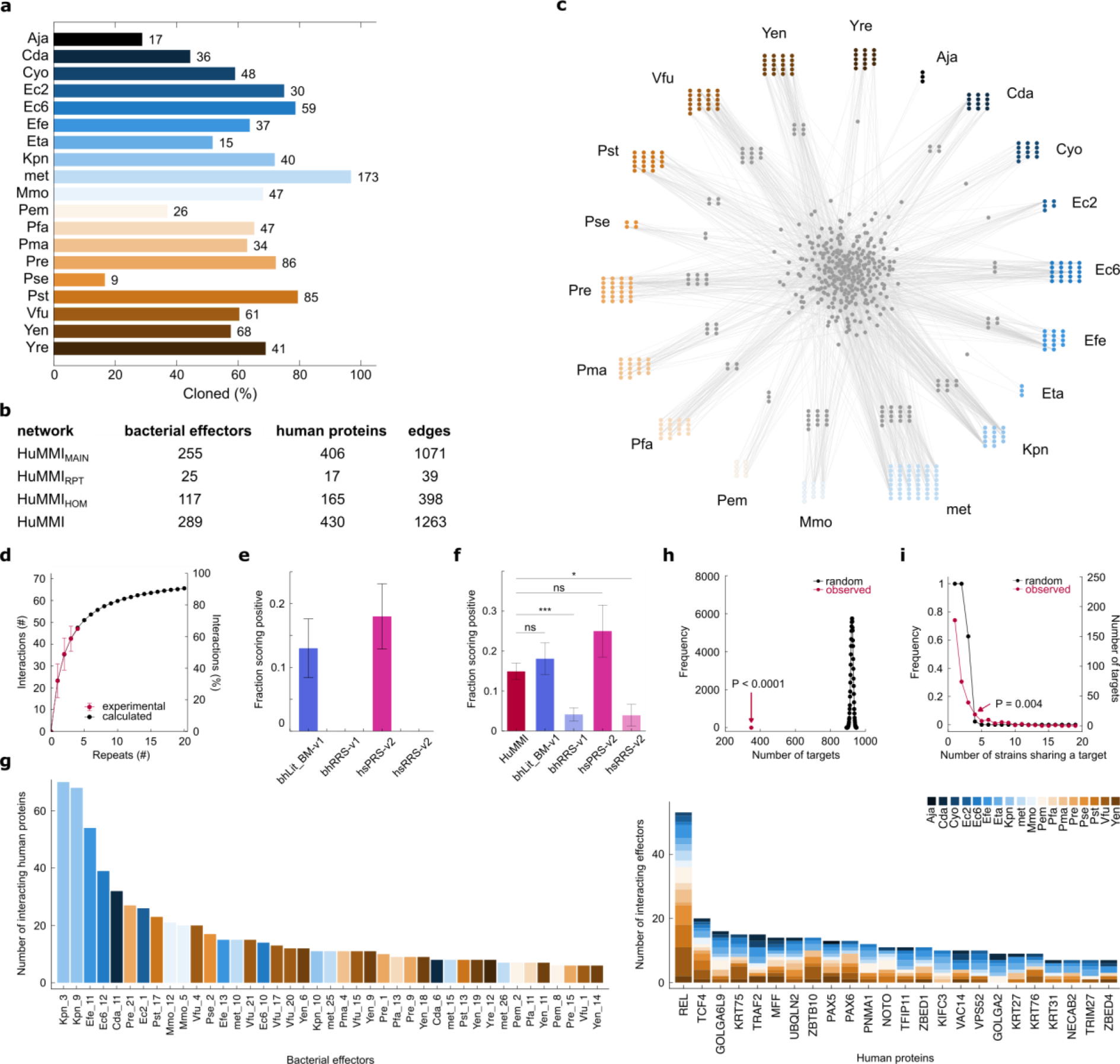
| Meta-interactome network map of bacterial effectors with human proteins. **a**, Success rates of effector ORF cloning for each strain, and number of sequence verified ORFs (right). **b**, Number of interactions and involved proteins in the HuMMI subsets. **c**, Verified human microbiome meta-interactome (HuMMI) map. Grey nodes: human proteins; outer layer human proteins targeted only by the nearest strain; central human proteins by effectors from multiple strains. **d**, Sampling sensitivity: saturation curve calculated from the repeat experiment: red dots represent average of verifiable interactions found in any combination of indicated number of repeat screens; black dots and line: modeled saturation curve. **e**, Assay sensitivity: percentage of identified interactions from bhLit_BM-v1 (n = 54 pairs), bhRRS-v1 (n = 73 pairs), hsPRS-v2 (n = 60 pairs), hrRRS-v2 (n = 78 pairs) in our Y2H. Error bars present the standard error (SE) of proportion. **f**, Validation rate of a random sample of HuMMI interactions (n = 295 pair configurations) compared to four reference sets in the yN2H validation assay: bhLit_BM-v1 (n = 94 pair configurations), bhRRS-v1 (n = 145 pair configurations), hsPRS-v2 (n = 44 pair configurations), hrRRS-v2 (n = 51 pair configurations). * *P* = 0.04; *** *P* = 0.0006; ns “no significant difference” (Fisher exact test; Extended Data Table 3). Error bars present SE of proportion. **g**, Left: degree distribution for the most connected effectors; right: effector-degree distribution for most targeted human proteins. Colors represent strains according to legend. **h**, Observed number of total effector targets in the human reference interactome (HuRI), compared to random expectation (exp. *P* < 0.0001; n = 10,000 randomizations). (**I**) Frequency distribution of human proteins targeted by effectors from the indicated number of different strains (red), compared to random expectation (black; n = 10,000). Targeting by effectors from four strains or more occurs significantly more often than expected by chance (exp. *P* = 0.004; n = 10,000).

To assess data quality, we assembled a positive control set of 67 well-documented manually curated binary interactions of bacterial (pathogen-) effectors with human proteins from the literature (bacterial human literature binary multiple – bhLit_BM-v1, Extended Data Table 3) and a corresponding negative control set of random bacterial and human protein pairs (bacterial host random reference set - bhRRS-v1). Benchmarking our Y2H assay in a single orientation with these and with the established human positive reference set (hsPRS-v2) and hsRRS-v2 indicated an assay sensitivity of ∼13% and 17.5%, respectively, which is consistent with previous observations^27,28^ (Fig. 2e and Extended Data Table 3). No negative control pair in either reference set scored positive, demonstrating the reliability of our system. In addition, we assessed the biophysical quality of HuMMI using the yeast nanoluciferase-2-hybrid assay (yN2H), which we benchmarked using the same four reference sets^25^. Notably, the retest rates of all sets involving bacterial proteins were lower than those of the human hsPRS-v2 and hsRRS-v2 across most of the scoring spectrum (Extended Data Fig. 2). Partly, this could be due to the nature of hsPRS-v2 pairs, which consist of very well-documented interaction pairs, which may have been selected for good detectability. In addition, the fact that the RRS sets exhibit the same overall trend indicates that interactions with prokaryotic proteins are more challenging to reproduce in this eukaryotic assay system, which reinforces the necessity for bacterial protein-specific reference sets (Fig. 2f, Extended Data Fig. 2, and Extended Data Table 3). At thresholds where the control sets were well separated, the retest rate of 173 randomly selected HuMMI interactions was statistically indistinguishable from the positive control sets, and significantly different from those of the negative controls (Fig. 2f, Extended Data Fig. 2, and Extended Data Table 3), indicating that the biophysical quality of our dataset is comparable to those of well-documented interactions in the curated literature.

The degree distribution of HuMMI_MAIN_ shows that numerous human proteins are targeted by multiple effectors (Fig. 2g and Extended Data Table 3), often from different species. Indeed, sampling analysis demonstrates that commensal effectors significantly converge on fewer host proteins than expected from a random process (Fig. 2h), thus suggesting selection for interactions with these targets. We had previously observed convergence of effectors from phylogenetically diverse pathogenic microbes on common proteins of their plant host^29,30^. In that system, we demonstrated with infection assays on genetic null mutant plant lines that the extent of convergence correlates with the importance of the respective host proteins for the outcome of the microbe-host interaction^29^. We therefore identified the human host proteins onto which commensal effectors converge. To this end, we sampled random effector targets for each strain and analyzed the distribution of repeatedly targeted proteins (Fig. 2i). While host proteins interacting with effectors from two strains are expected at high frequency by chance, targeting by four bacterial strains is unlikely to emerge by chance (Fig. 2i and Extended Data Table 3). Thus, the 60 human proteins targeted by effectors from four or even more commensal strains are subject to effector convergence and may be of general importance for human microbe-host interactions. Together with our recently published plant-symbiont interaction data^31^, these data suggest that convergence has evolved as a universal feature of effector-host interactions independent of the microbial lifestyle and kingdom of the host organism.

### Sequence features mediating effector-host interactions

The function of unknown proteins can often be inferred from better-studied orthologues, but convergence could also result from high sequence similarity among effectors. We therefore compared sequence- to interaction-similarity as a proxy for their function in host cells (Fig. 3a). Within the systematically retested HuMMI_HOM_ clusters, both are poorly correlated and sequence similarity merely defines the upper limit for interaction similarity but does not imply it. This is illustrated by cluster 3, in which all seven effectors share over 90% mutual sequence similarity while their pairwise interaction profile similarities range from identical to complementary (Fig. 3b and Extended Data Table 3).

**Fig. 3.**
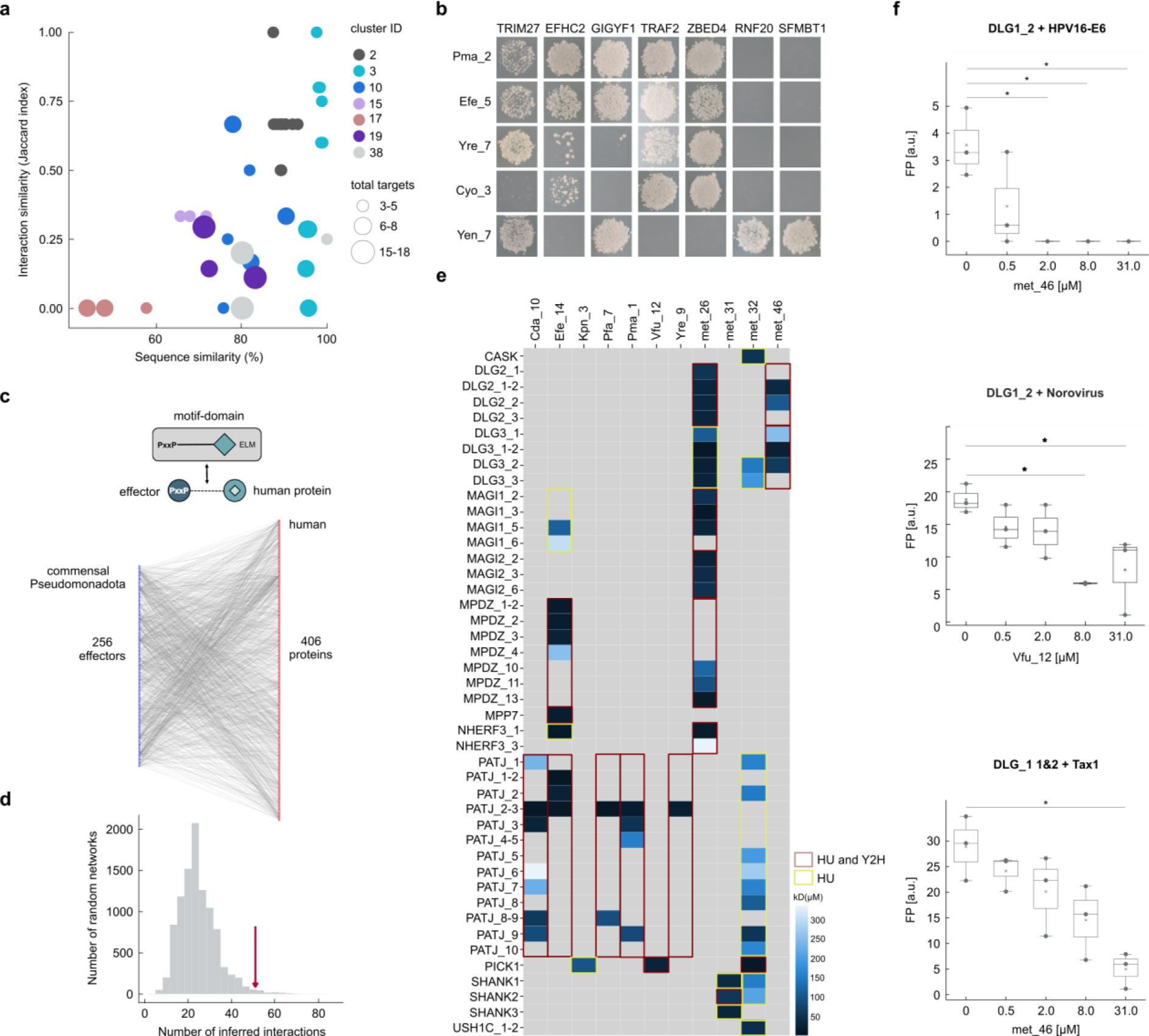
| Interaction specificity and interaction motifs. **a**, Scatter plot of sequence- and Jaccard-interaction similarity for all effector pairs within indicated homology groups of HuMMI_HOM_ with ≥ 3 interactors and effectors. Node size indicates union of human proteins targeted by effector-pair according to legend. **b**, Y2H data for one of four repeats for homology cluster 3. **c**, Schematic of interaction motif-domain interface identification in the effector-host interaction. **d**, Count of motif-domain pairs matching at least one stringency criteria identified in HuMMI_MAIN_ (arrow) compared to random expectation (experimental *P* value, n = 10,000). **e**, Interaction strength of PDZ domains of human proteins with C-terminal 10 amino acid peptides of the effectors indicated on top. Calculated *K_D_* according to legend. Overlap between HU and Y2H is indicated by colored frames. **f**, Competition of the interaction between human PDZ domains and viral PBM peptides by the indicated effector peptides. * *P* < 0.05 (Kruskal Wallis with Dunn’s correction, n = 3). Boxes represent interquartile range (IQR), with the bold black line representing mean; whiskers indicate highest and lowest data point within 1.5 IQR.

Using HuMMI_MAIN_ we also investigated if effectors without substantial sequence similarity share interaction similarity, which might indicate shared functions. In fact, clustering effectors by their pairwise interaction similarity identified substantial overlap outside the homology clusters (Extended Data Fig. 3), indicating that dissimilar effectors may have similar functions in the host. Both analyses indicate that effector function as measured by protein-interaction profiles is largely independent of overall sequence similarity.

Looking for structural correlates for interaction specificity, we wondered whether domain-domain or domain-short linear motif (SLiM) interfaces mediating the interactions can be identified (Fig. 3c). Using experimentally identified interaction templates^32^, a putative interface was found for 52 interactions in the HuMMI_MAIN_ screen (Extended Data Table 4). Of these, 43 interactions matched motif-domain templates passing one (Fig. 3d), and 22 passing two stringency criteria (Extended Data Fig. 3). Among the former, 23 interactions involve PDZ domains in the human protein, which recognize PDZ-binding motifs (PBM) in the C-terminus of interacting bacterial proteins. PDZ domain-containing proteins commonly mediate cell-cell adhesion, cellular protein trafficking, tissue integrity, as well as neuronal and immune signaling^33^. To experimentally validate these interfaces, individual and tandem PDZ domains from 13 human proteins and C-terminal peptides from 16 interacting bacterial effectors were tested via Holdup, a quantitative chromatographic *in vitro* interaction assay^34,35^. For 16 of 23 Y2H pairs (70%) at least one PDZ-peptide interaction was identified, all with affinities between 1 and 200 µM (Fig. 3e and Extended Data Table 4). In three instances two PDZ domains arranged in tandem were required to detect the interaction by Holdup, indicating that some Y2H pairs might have been missed because not all PDZ combinations of the proteins were tested. For human proteins with multiple PDZ domains, often different domains were the target for different effectors demonstrating both specificity and functional specialization of the effectors (Fig. 3e).

Because of their functioning in immune signaling and cell shape, PDZ domains are frequently targeted by viruses^36^. This opens the possibility that bacterial effectors and viral proteins compete for PDZ-binding and thus mutually influence their respective impact on the host. To gather support for this possibility, we identified viruses that can cause infections in the digestive tract, namely SARS-CoV-2^37^, HPV16 and 18, which have a high prevalence in human guts and have been linked to colorectal cancer^38^, and norovirus, a globally common cause of gastroenteritis and diarrhea^39^. We selected two hitherto unpublished interactions of Norovirus VP2 C-terminal peptide with DLG1 (domain 2) and MAGI1 (domain 4), and previously observed interactions between the C-terminal peptides of SARS-CoV-2 E with SHANK3, and of HPV16 and 18 E6 with the PDZ domains of PICK1 and MAGI4 (domain 1), respectively^34^. Indeed, in fluorescent polarization assays the viral PBM peptides competed with those of the effectors Vfu_12, met_32, met_31, and met_46 (Fig. 3f and Extended Data Fig. 3). Similarly, the functionally well-characterized interaction of the C-terminus of HTLV1 Tax1 with DLG1^40^ was competed off by the met_32 PBM peptide. Thus, viral and bacterial proteins may compete in the intracellular environment for binding partners and hence for influence on human cell function. Such competition could contribute to the previously observed mutual influence of microbiome and viral infection on each other^41^.

Thus, while the overall sequence similarity of effectors does not correlate with their host-protein interaction profiles, several interfaces mediating the interactions can be identified. How these interactions compete with human and viral proteins to modulate the host network is an important question for future studies.

### Effector-targeted functions and disease modules

To explore the potential roles of commensal effectors in the host we analyzed the functions of the targeted human proteins through gene ontology (GO) enrichment analysis (Fig. 4a, Extended Data Fig. 4, and Extended Data Table 5). Redundant parent-child GO-term pairs were grouped and are displayed by a representative term. Intriguingly, “response to muramyl-dipeptide (MDP)”, a bacterial cell wall-derived peptide that can be perceived by human cells, was among the most enriched functions, thus not only supporting the relevance of our interactions but indicating that effectors modulate cellular responses to their detection. Moreover, a key component of the MDP signaling pathway is NOD2, which is encoded by a major susceptibility gene for Crohn’s disease (CD)^42^, an autoimmune disease with a strong etiological microbiome contribution^43^. In addition, several central immune signaling pathways are enriched among the targets, namely the NF-κB and the stress-activated protein kinase and Jun-N-terminal kinase (SAPK/JNK) pathways, supporting the notion that modulation of immune signaling is an important function of commensal effectors. Remarkably, five of the significantly targeted convergence-proteins belong to the NF-κB module (Extended Data Fig. 4), one of the evolutionarily oldest immune signaling pathways in animals that is already present in sponges^44^. This may reflect the long co-evolution between microbial effectors and this ancient immune coordinator. Relating to human disease, anti-TNF biologicals, which dampen NF-κB-driven immunity, are an important therapeutic for diverse autoimmune diseases including CD, psoriasis, and rheumatoid arthritis. Another highly enriched group of five terms relates to collagen production, which suggests that effectors may modulate the extracellular environment that hosts the microbes. Inflammation-independent fibrotic collagen production is an important clinical feature of CD, and the gut microbiota has been found to be a main driver^45^. As several metabolism-related terms were identified, we also tested directly whether enzymes in the Recon3D^46^ model of human metabolism were targeted. Indeed, we detected a significant enrichment of metabolic enzymes (*P* = 0.0001, Fisher’s exact test) and nominally significant targeting of bile acid and glycerophospholipid metabolism, and fatty acid oxidation (Extended Data Table 5). Overall, however, despite the strong overall signal and general targeting of fatty acid metabolism, no individual metabolic subsystem stood out as being targeted by effectors from more than two strains or having more than two targeted proteins.

**Fig. 4.**
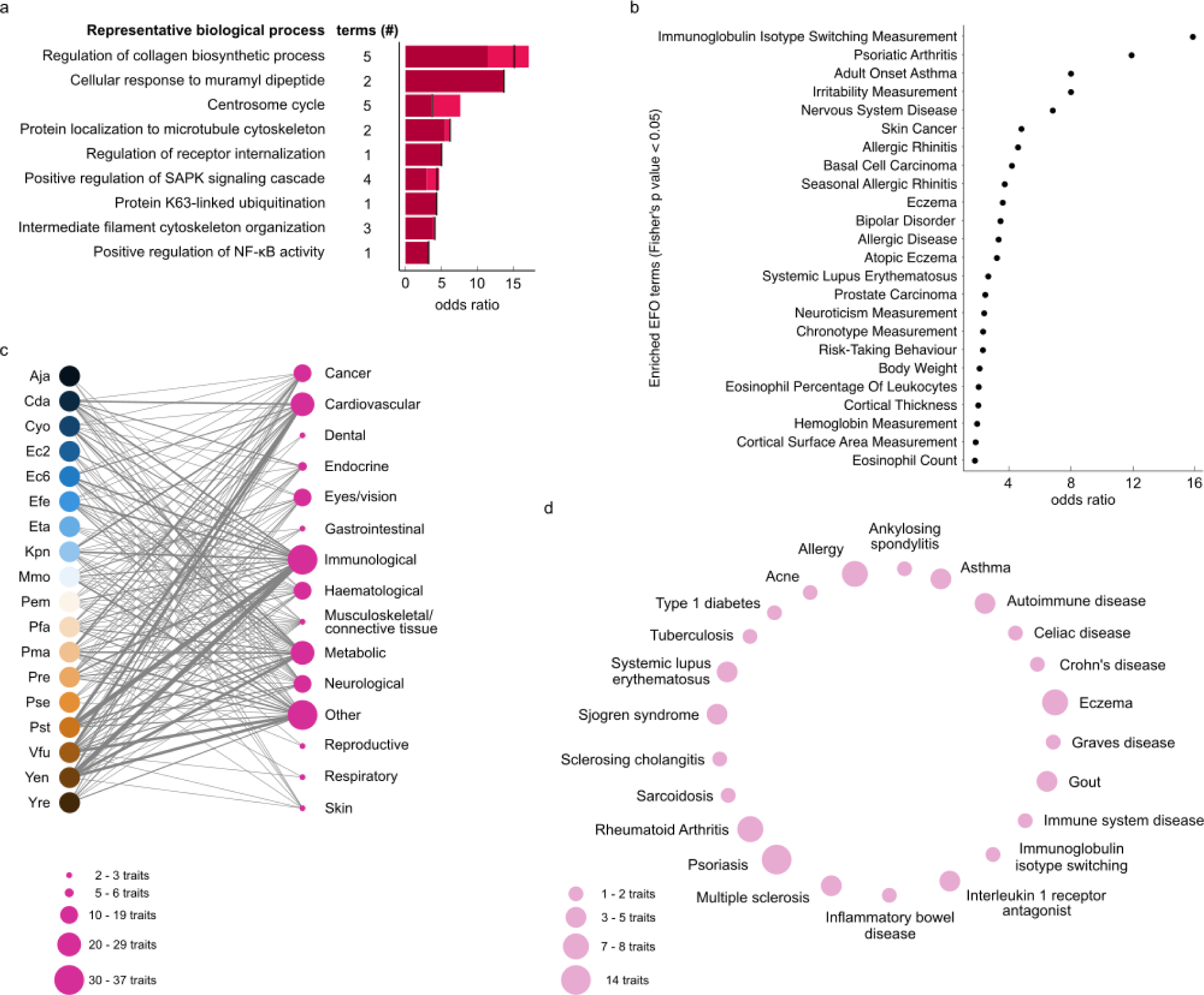
| Function and disease association of microbially targeted human proteins. **a**, Odds ratios (OR) of representative functional annotations enriched among effector targeted human proteins (FDR < 0,05, Fisher’s exact test with Bonferroni FDR correction). The number of represented terms is shown by terms (#). The lowest and highest OR observed for the represented group are indicated by light shaded area in each bar. Black line indicates OR for representative term. Full data and precise FDR and OR values in Extended Data Table 5. **b**, Genetic predisposition for traits and diseases enriched among human genes encoding effector targets in HuRI (cutoff FDR = 0.05, Fisher’s exact test, n = 349). **c**, Disease groups for which genetic predisposition is enriched in network neighborhoods of effectors from the indicated strains. Trait node size corresponds to number of significantly targeted traits in that group according to legend. Stroke of strain-group edge reflects number of underlying significant effector-trait links (α < 0.01 and OR > 3, Fisher’s exact test). **d**, Specific diseases underlying the ‘immunological’ group in c. Node size reflects the number of underlying effector-trait associations according to legend.

From a network perspective, proteins encoded by disease-genes (disease proteins) constitute nodes and form disease modules^47^, whose functional perturbation promotes pathogenesis. Importantly, viruses can contribute to non-infectious disease etiology by binding to and similarly perturbing these disease proteins and modules^48^. Therefore, we wondered if bacterial effectors also target such network elements and may thereby influence human traits. We started with “causal genes/proteins” identified from genome-wide-association studies (GWAS) by the Open Targets initiative^49^, and merged gene sets for traits identified as identical by their experimental factor ontology (EFO) terms (Extended Data Table 5). We first investigated direct effector targets. The strong enrichment of the “immunoglobulin isotype switching” trait among these is intriguing as the evolutionarily older IgA antibodies are emerging as having an important role in shaping the gut microbiome^50,51^. Effector-targeted proteins are further associated with diverse cancers and with diseases that have a strong immunological component, including asthma, psoriasis, allergies, and systemic lupus erythematosus (Fig. 4b, cutoff nominal *P* = 0.05, Fisher’s exact test, Extended Data Table 5). While none of the identified diseases is currently known as an ailment of the gut it has emerged that the gut microbiome shapes immune homeostasis and contributes to lung and skin diseases like asthma^52^ and psoriasis^53^. In addition, some of the disease-associated genes encode convergence proteins for effectors from multiple bacterial species (Fig. 2g). As such, it is plausible that proteins like REL or TCF4 are similarly targeted by effectors from Pseudomonadota in skin or lung microbiome communities and contribute to the identified diseases. Moreover, 26% of the effectors in HuMMI are also detectable in skin microbiome samples (Extended Data Table 5), indicating that commensal effectors are shared between different ecological niches.

A partly complementary explanation emerges from our previous studies of human and plant pathogen-host systems. In these evolutionary distant systems, we showed that genetic variation affecting the severity of infection does not reside in genes encoding direct targets but in interacting, i.e., neighboring proteins in the host network^25,29^. We, therefore, explored the network neighborhood of all effector-targets using short random walks in the human reference interactome (HuRI)^54^. We identified proteins that were significantly more often visited in HuRI compared to degree-preserved randomly rewired networks, which we considered the ‘neighborhood’. For each effector-targeted neighborhood, we assessed the enrichment of gene products associated with diverse human traits using Open Targets causal genes. Nominally significant associations were aggregated on a strain level and summarized for disease groups (Fig. 4c and Extended Data Table 5). Intriguingly, most disease groups for which susceptibility-gene products are enriched in the target neighborhoods represent traits that have been linked to the gut microbiome^55^. Apart from immunological traits, these include cardiovascular, metabolic, and neurological traits as well as multiple cancers, including colorectal cancer. Among the target neighborhoods for immunological diseases, we identified associations to CD (nominal *P* = 8.5 * 10^-5^, Fisher’s exact test) and inflammatory bowel disease (nominal *P* = 0.0008, Fisher’s exact test) but not to ulcerative colitis (UC) (Fig. 4d and Extended Data Table 5). Neighborhoods harboring genetic susceptibility associated with psoriatic arthritis, asthma, and allergies were also significantly targeted, which recapitulates the observations for direct targets. Considering the importance of the microbiome for human metabolic disorders^55^ it is noteworthy that network modules important for HDL and LDL cholesterol levels (nominal *P* = 0.006 and *P* = 0.008, respectively, Fisher’s exact test), and several diabetes traits were significantly targeted albeit less recurrently than inflammatory diseases and cancers (Extended Data Table 5). Together, these results suggest that commensal effectors modulate their host’s immune system and local metabolic and structural microenvironment. As genetic variation affecting the targeted proteins and their network neighborhood is linked to several human diseases, functional modulation of the same network neighborhoods by commensal effectors may contribute to disease etiology. The fact that the risk for several of the identified diseases is known to be modulated by the microbiome strengthens this hypothesis. We therefore investigated if commensal effectors, indeed, perturb some of the identified pathways and functions.

### Effector function in human cells and disease

The NF-κB signaling module is enriched among the convergence proteins and all targets of commensal effectors (Fig. 4a and Extended Data Fig. 4). Because of its important role in many diseases, we chose a cell-based dual-luciferase assay^25^ to test whether commensal effectors modulate NF-κB pathway activity in human cells. Indeed, five of 26 commensal effectors caused a significant increase in NF-κB pathway activity in the absence of exogenous stimulation suggesting pathway activation (Fig. 5a and Extended Data Table 6). Conversely, three effectors significantly reduced relative transcriptional NF-κB activity even in the presence of strong TNF stimulation (Fig. 5b, Extended Data Fig. 5, and Extended Data Table 6). Since some bacterial effectors also modulate NF-κB-independent induction of the thymidine kinase control promoter, we assessed the impact of selected effectors on endogenous expression of NF-κB controlled human adhesion factor ICAM1 and cytokine secretion. We focused these experiments on two NF-κB activating (Kpn_9, met_7) and two NF-κB inhibiting (Pst_11, Cyo_12) bacterial effectors. ICAM1/CD54 is a glycoprotein that mediates intercellular epithelial adhesion and interactions with immune cells, specifically neutrophils. Epidemiologically, ICAM1 has been linked to CD such that increased ICAM1 expression is associated with higher disease risk^56^ likely by facilitating recruitment and retention of inflammatory immune cells^57,58^. Interference with ICAM1-mediated neutrophil trafficking is currently being tested as a therapeutic approach to treat CD^59^. In colon carcinoma Caco-2 cells, expression of met_7 caused a significant increase of ICAM1 expression (*P* = 0.05, one-way ANOVA with Dunnett’s multiple hypothesis correction, Extended Data Table 6) following stimulation with a pro-inflammatory cocktail. Expression of the inhibitory effectors Pst_11 and Cyo_12 did not significantly alter the induction of ICAM1 cell surface expression (Fig. 5c). We also investigated the effect of met_7 and Cyo_12 on cytokine secretion in unstimulated Caco-2 cells or following pro-inflammatory stimulation. In basal conditions, Cyo_12 reduced the secretion of several cytokines especially IL6 and IL8, whereas met_7 caused an increase in IL8 secretion in these conditions (Fig. 5d and Extended Data Table 6). Following proinflammatory stimulation, expression of Cyo_12 further reduced cytokine secretion. This effect was most pronounced for IL8, but also significant for IL6 and the pro-inflammatory IL1beta, IL18, and IL23. These cytokines are noteworthy as they are linked to IBD pathogenesis. IL23R has been associated to CD, and IL6 and IL23 stimulate the differentiation of Th17 cells, which have emerged as key players in CD^60,61^. IL8 is overexpressed in colonic tissue of IBD patients and has been suggested as a chemoattractant triggering neutrophil invasion^62,63^. In contrast, no significant impact of met_7 on cytokine secretion was detectable in the context of stimulation (Fig. 5e and Extended Data Fig. 5). Thus, commensal effectors can both stimulate and dampen intracellular immune signaling and this modulation can impact immune and tissue homeostasis via cell-cell adhesion and cytokine secretion.

**Fig. 5.**
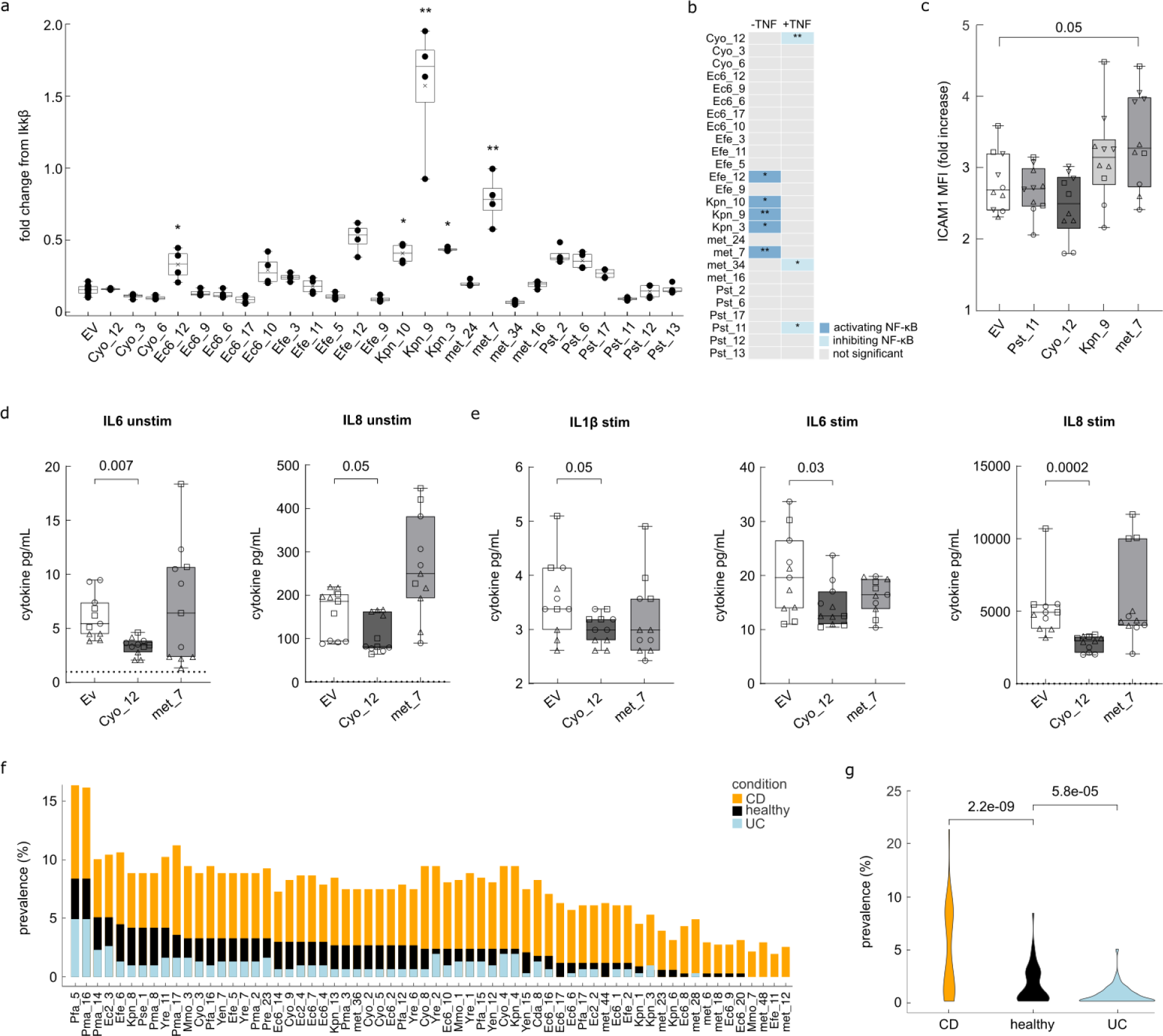
| Effector impact on human cell function and clinical prevalence in IBDs. **a**, Relative NF-κB transcriptional reporter activity of HEK293 cells expressing the indicated effectors or empty vector (EV) in unstimulated conditions (Kruskal-Wallis test with Dunn’s correction, * *P* < 0.05, ** *P* < 0.01, n = 4). Boxes represent IQR, black line indicates the mean, whiskers indicate highest and lowest data point within 1.5 IQR. **b**, Summary of significant impact of effectors on normalized NF-κB transcriptional reporter activity in baseline conditions and after TNF stimulation (Kruskal-Wallis test with Dunn’s correction, * *P* < 0.05, ** *P* < 0.01, n = 4). **c**, Fold-induction of ICAM1 expression following pro-inflammatory stimulation of Caco-2 cells transfected with the indicated effectors (one-way ANOVA with Dunnett’s multiple comparison test, n = 10). **d**, Concentration of cytokines secreted by Caco-2 cells in basal conditions transfected with the indicated effectors. EV indicates empty vector mock control. *P* values calculated by Kruskal-Wallis test (n = 11). Dashed line indicates detection limit of assay. **e**, Concentration of cytokines secreted by Caco-2 cells stimulated by a pro-inflammatory cocktail transfected with the indicated effectors. EV indicates empty vector mock control. Indicated *P* values calculated by Kruskal-Wallis test (n = 11). Dashed line: detection limit of assay. C – E Boxes represent IQR, black line indicates the mean, whiskers indicate highest and lowest data point. **f**, Effector prevalence in metagenomes of CD (n = 504), and UC patients (n = 302) compared to healthy controls. Effectors are significantly more prevalent in CD patient metagenomes (FDR < 0.01; Fisher exact test, Benjamini-Hochberg correction). **g**, Effector prevalence distribution among the indicated cohorts. *P* values calculated by Wilcoxon rank- sum test, n as in f.

As we identified both genetic and functional links between commensal effectors and IBD-related processes, we sought clinical evidence for a potential role of effectors in these diseases. We hypothesized that a potential role of effectors in IBD etiology may be reflected in altered effector prevalence in the microbiota of patients versus healthy controls. Analyzing a large dataset with > 800 IBD patient-derived and > 300 healthy control-derived metagenomes^64^ we found 64 effectors that were significantly more prevalent in the metagenomes of CD patients compared to healthy controls (Fig. 5f and Extended Data Table 6). In metagenomes of UC patients only three effectors had a significantly different prevalence, and, intriguingly, these were less common compared to healthy controls (Extended Data Table 6). This trend was recapitulated when the prevalence distributions of all detected effectors were analyzed. Whereas CD patients had a significantly higher load of effectors, the overall effector prevalence was lower in UC patients compared to healthy subjects (Fig. 5g and Extended Data Table 6). These opposing findings were unexpected as an increased abundance of Pseudomonadota has been reported both for CD and UC patients^65^. At the same time, many clinical features such as affected tissues and response to anti-TNF therapy differ between these two forms of IBD, rendering it plausible that effectors contribute differently to their etiology. Whether commensal effectors indeed causally contribute to disease etiology or acute flairs is an important question with potential therapeutic implications.

## Discussion

The presence of T3SS in human commensal microbes has been noticed previously and was speculated to mediate crosstalk between the intestinal microbiota and the human host^66,67^. Here, we provide evidence that, analogous to the plant kingdom^31,68^, also in the human gut T3SS and effectors function in commensal microbe-host interactions and modulate immune signaling. Thus, effector secretion appears to be used universally by Pseudomonadota to mediate interactions with multicellular eukaryotes independently of the lifestyle of the microbe.

Since, as we show, commensal effectors modulate immune signaling we hypothesized that this may affect the manifestation of human diseases, especially those involving the immune system. The influence of the microbiome on IBD etiology is well documented^1^. Therefore, it is noteworthy that IBD, especially CD, emerged in several of our analyses. Effectors target the “response to the muramyl-dipeptide” pathway which includes NOD2, a major CD-associated gene product^69^. Further, effectors target and regulate the NF-κB pathway, which is strongly activated by TNF, a key therapeutic target in CD^70^. Likewise, ICAM1 is a susceptibility gene for CD whereby high expression increases disease risk^56^. Secretion of IL6, IL8, and IL23 is significantly altered by effectors, and all have previously been linked to CD^61,63^. Thus, commensal effectors regulate several IBD-relevant pathways and can thus influence the establishment or maintenance of feedback loops during disease development^71^. This conclusion is strengthened by the observation that effectors are enriched in metagenomes of a CD patient cohort. Thus, multiple lines of evidence suggest that by modulating immune signaling, commensal effectors contribute to the etiology of CD.

Likely other microbial habitats of the human body, such as skin or lung, also host T3SS+ strains, and we identified effectors in a skin metagenome. It will be important to investigate this in the future to understand if those effectors have similar targets and effects on local cells. ICAM1, e.g., is the entry receptor for rhinovirus A^72^, and an increased expression due to microbial effectors could increase the risk for infections and thus to develop asthma^73,74^. The broader question of how effectors influence the pathogenesis of IBD and other diseases will be important to address in further detailed studies. Our molecular data show that different effectors can have opposing impacts on immune pathways, analogous to genetic variants. Thus, host genetics and effectors jointly impact on the molecular networks, and pathogenic developments emerge from the interplay of protective and disease enhancing factors. For CD specifically, however, our analyses suggest that effectors promote disease development.

In summary, we demonstrate that bacterial effector proteins constitute a hitherto unrecognized regulatory layer by which the commensal microbiota communicates with host cells and modulates human physiology. We anticipate that our findings and resources will open new research directions towards understanding the host-genetics dependent mechanisms by which the microbiome influences human health and exploring the potential of effectors for therapy and prevention.

## METHODS

### Identification of T3SS+ strains in culture collections and MAGs

To collect reference genomes for strains available from culture collections, three large culture collections were queried for all Pseudomonadota strains: DSMZ via BacDive^75^, ATCC (atcc.org) and BEI (beiresources.org). The strain numbers were looked up in GenBank (Release 229) from which 77 strains could be identified as perfect match.

MAGs that were at least 50% complete and less than 5% contaminated (as estimated by CheckM^76^ from two different meta-studies were selected. 92,143 MAGs of Almeida et al.^15^ and 9,367 Pseudomonadota MAGs from Pasolli et al.^16^ were used as input for T3SS prediction scaled via massive parallel computing. The computational predictions presented have been achieved in part using the Vienna Scientific Cluster (VSC). The prediction performance of EffectiveDB^10^ on incomplete and contaminated MAGs was assessed by 5-fold cross-validation with 5 repeats using 0 - 100% completeness and 0 - 50% contamination in 5% steps of simulated incompleteness/contamination, randomly sampling genes from test-set. In addition, T3SS were predicted for 4,753 strains isolated by the human gastrointestinal bacteria genome collection (HBC)^11^, and the unified gastrointestinal genome (UHGG) collection^12,13^. A performance-improved re-implementation of the EffectiveDB classifier (https://github.com/univieCUBE/phenotrex, trained on EggNOG 4 annotations^77^) was used to predict functional T3SS present in MAGs and genomes of isolated strains. Threshold for positive prediction was defined as > 0.7.

Protein sequences were predicted from 44 T3SS-positive reference strains and MAGs using prodigal v2.6.3^76^. Of 770 MAGs a total of 474,871 representative protein sequences were identified using CD-HIT^78^ (v4.8.1, parameters: ‘-c 1.0’). The identical procedure was performed for 44 genomes from culture collections resulting in 161,115 proteins. Machine-learning based tools were used to predict T3SS signals (EffectiveT3 v.2.0.1 and DeepT3 2.0^19^) or effector homology using pEffect^21^ to extract potential effector proteins. The results of all three tools were combined using a 0 - 2 scoring scheme: 2 for perfect score (pEffect > 90, EffectiveT3 > 0.9999, DeepT3: both classifiers positive prediction), 1 for positive prediction as defined by default settings (pEffect > 50, EffetiveT3 > 0.95, DeepT3: one classifier) and 0 for negative prediction. Sequences with a sum score above 4 were regarded as potential effectors. Further, all sequences without start/stop-codon or trans-membrane region containing proteins (> 0 regions; predicted with TMHMM version 2.0) were excluded. Proteins were clustered using 90% sequence identity threshold (CD-HIT parameters ‘-c 0.9 -s 0.9’) to reduce redundancy. Effector-clusters with great diversity regarding T3SE-prediction scores were removed from the final set. Full data in Extended Data Table 1.

### Identification of effector similarities and homology groups

Based on a mutual sequence identity of ≥ 30% over 90% of the common sequence length effectors were considered ‘homologous’ and included in the HuMMI_HOM_ experiment to investigate the impact of sequence similarity on interaction similarity. Protein sequences were analyzed by global alignment using Needleman Wunsch algorithm implemented in the emboss package (Extended Data Table 2).

### Commensal vs pathogen effector similarity

We gathered the sequences of 1,195 known pathogenic T3 effectors from the BastionHub database^79^ (August 29^th^, 2022). We assessed the similarity between commensal and pathogenic effector sequences using BLAST (stand-alone, version 2.10^80^). For each commensal effector, the pathogen effector with the highest sequence similarity was considered as best match. Subsequently, we computed the alignment coverage over the pathogenic effector sequence. Full data in Extended Data Table 2.

### Cohort analyses

Genomes of bacterial isolates from the human gut were gathered from multiple published datasets^11–13^. The presence of T3SS was predicted for each of these genomes as described above. GTDB-Tk (v2.1)^81^ was used to assign the taxonomy to each of the genomes, and the concatenated bac120 marker proteins from this were used to generate a phylogenomic tree of the isolates, visualized in iTOL^82^. FastANI was used to match the T3SS positive genomes to the WIS representative genomes of the human gut^18^ based on ANI values > 95%^83^. The relative abundance of the 10 matching representative genomes was then identified across 3,096 Israeli, and 1,528 Dutch patients^18^.

### Effector cloning

Bacterial strains from the ATCC collection were ordered from LGS Standard Standard (Wesel, Germany) or ATCC in the US (Manassas, Virginia). Bacterial strains from the DSMZ collection were obtained from the Leibniz-Institut DSMZ (Braunschweig, Germany) and strains from the BEI collection were ordered at BEI resources (Manassas, Virginia, USA) (Extended Data Table 2). Effectors identified from MAGs and effectors for the PRS were ordered at Twist Bioscience (San Francisco, CA, 660 USA). If no genomic DNA could be obtained strains were cultured according to the manufacturer’s protocol and DNA was extracted using the NucleosSpin Plasmid (NoLid) Mini kit (Macherey-Nagel cat. No. 740499) with vortexing after addition of BufferA2 and BufferA3. A nested PCR was performed to add Sfi sites, the DNA was purified using magnetic beads (magtivio cat. no. MDKT00010075), followed by an Sfi digestion and another clean-up with magnetic beads. Digested PCR products were cloned into pENTR223.1 using T4 DNA Ligase (ThermoFisher ca. no. EL0011). Plasmids were propagated in DH5α *E. coli* and the plasmid DNA was extracted using the pipetting Bio Robot Universal System (Qiagen cat. no. 9001094) and the QIAprep 96 plus BioRobot kit (Qiagen cat. no. 962241). ORFs were verified by Sanger Sequencing. Effectors were cloned into the Y2H destination plasmid pDEST-DB (pPC97, Cen origin), the pDEST-N2H-N1 and -N2, or the mammalian expression vector pMH-FLAG-HA by an LR reaction of the Gateway System. After propagation in DH5α *E. coli* and DNA extraction plasmids were transformed into *S. cerevisiae* Y8930 (MATα mating type) as DB-X ORFs as described^84^.

### Meta-interactome mapping

A state-of-the-art high-quality Y2H screening pipeline was followed as previously described^25,85^. DB-X ORFs were tested for autoactivation by mating against AD-empty plasmids in Y8800 (MATa). 45 ORFs of the strains and 14 meta effectors tested positive and were excluded from subsequent steps. The remaining 900 ORFs were individually mated against pools of ∼188 AD-Y human ORFs from the human ORFeome collection v9.1 including 17,472 ORFs^86^. During primary screening, haploid AD-Y and DB-X yeast cultures were spotted on top of each other and grown on yeast extract peptone dextrose (YEPD) agar (1%) plates. After incubation for 24 h, the clones were replica plated onto selective synthetic complete media lacking leucine, tryptophan and histidine (SC-Leu-Trp-His) + 1 mM 3-AT (3-amino-1,2,4-triazole) (3-AT plates) and replica cleaned after 24 h. 48 h later, three colonies were picked per spot and grown for 72h in SC-Leu-Trp liquid medium. For the secondary phenotyping, yeasts were spotted on SC-Leu-Trp plates and after incubation for 48 h replica plated and cleaned on 3-AT-plates and SC-Leu-His + 1 mM 3-AT + 1 mg per litre cycloheximide plates to identify spontaneous DB-X autoactivators. Clones growing on 3-AT plates, but not on cycloheximide plates were picked into yeast lysis and processed to generate a library for pair identification by Next Generation Sequencing using a modified KiloSeq procedure as previously described^25^. Identified DB-X and AD-Y pairs were mated individually during the fourfold verification, replica plated and cleaned after 24 hours and picked after another 48 h incubation. Growth scoring was performed using a custom dilated convolutional neural network as described^25^. Pairs scoring positive at least three out of the four repeats qualified as bona fide Y2H interactors. The AD-Y and DB-X constructs were identified once more by NGS. All interaction data are in Extended Data Table 3.

### Assembling reference sets

To identify additional reliably documented interactions between bacterial effectors and human proteins for the positive control set (bhLit_BM-v1), we queried the IMEx consortium protein interaction databases^87^ through the PSICQUIC webservice^88^ (May 10^th^, 2021) using the T3 effectors UniprotKB accession numbers and fetched all the PubMed identifiers of the articles describing additional interactions. In total, we gathered 67 interactions between 29 T3 effectors and 64 human proteins, described in 13 distinct publications that underwent the manual curation step for inclusion in the PRS (Extended Data Table 3).

### Y2H assay sensitivity

Effector ORFs from bhLit_BM-v1 and bhRRS-v1 (Extended Data Table 3) were transferred into pDEST-DB (DB-X) and transformed into *Saccharomyces cerevisiae* Y8930 (MATα). Yeast strains containing the corresponding AD-Y human ORF were picked from hORFeome9.1^86^ and ORF identity verified by end-read Sanger sequencing of PCR products. Yeast strains harboring plasmids containing ORFs from hsPRS-v2/hsRRS-v2^89^ were provided by the Center for Cancer Systems Biology, Dana-Farber Cancer Institute, Boston, MA. DB-X and AD-Y were mated fourfold with each other, as well as against yeast strains containing the corresponding DB-empty or AD-empty plasmid. Growth scoring was performed as described above for the fourfold verification. Pairs scoring positive at least three out of the four repeats qualified as bona fide Y2H interactors.

### Interactome validation by yN2H

200 interactions were randomly picked from HuMMI and all ORFs from the indicated datasets (Extended Data Table 3) were transferred by Gateway LR reactions into pDEST-N2H-N1 and pDEST-N2H-N2 plasmids containing a *LEU2* or *TRP1* auxotrophy marker, respectively^89^. Successful cloning was monitored by PCR-mediated evaluation of insert size, and positive clones transformed into haploid *Saccharomyces cerevisiae* Y8930 (MATα) and Y8800 (MATa) strains, respectively. Protein pairs from all datasets were randomly distributed across matching 96-well plates.

5 µL of each haploid culture of opposite mating type grown to saturation was mated in 160 µL YEPD medium and incubated overnight. Additionally, each position was mated with yeast stains containing empty N1 or N2 plasmids, to measure background. 10 µL mated culture was inoculated in 160 µL SC-Leu-Trp and grown overnight. 50 µL of this overnight culture was reinoculated in 1.2 ml SC-Leu-Trp and incubated for 24 h at 1000 rpm. Cells were harvested 15 min at 3000 rpm, the supernatant discarded, and each cell pellet was fully resuspended in 100 µl NanoLuc Assay solution (Promega corp. Madison, WI, USA, cat# 1120). Homogenized solutions were transferred to white flat-bottom 96-well plates (Greiner Bio-One, Frickenhausen, Germany, cat# 655904) and incubated in the dark for 1 h at room temperature. Luminescence for each sample was measured on a SpectraMax ID3 (Molecular Devices, San Jose, CA, USA) with 2 s integration time. The normalized luminescence ratio (NLR) was calculated by dividing the raw luminescence of each pair (N1-X N2-Y) by the maximum luminescence value of one of the two background measurements. All obtained NLR values were log_2_ transformed and the positive fraction for each dataset was determined at log_2_ NLR thresholds between –2 and 2, in 0.01 increments. Statistical results were robust across a wide range of stringency thresholds. Extended Data Table 3 reports the results at log_2_ NLR = 0. Reported *P* values were calculated by Fisher’s exact test.

### Interactome framework parameter calculation

*Assay sensitivity* (S_a_), i.e., the fraction of detectable interactions was assessed employing the effector bhLit_BM-v1 (54 pairs) and bhRRS-v1 (73 pairs) as well as the human hsPRS-v2 (60 pairs) and hsRRS-v2 (78 pairs) for benchmarking. All reference sets were tested 4 times using the Y2H screening pipeline. To assess sampling sensitivity (S_s_) a repeat screen was conducted. 288 bacterial effectors were screened 4 times against 5 pools comprising 1,475 human proteins. A saturation curve was calculated as described^85^. Briefly, all combinations of the number of interactions of the 4 repeats were assembled and the reciprocal values calculated. From these a linear regression was determined to obtain the slope and the intercept. Reciprocal parameters were calculated to find V_max_ and K_m_ and using the Michaelis-Menten-formula a saturation curve was predicted. *Overall sensitivity* emerges from both sampling and assay limitations and is calculated as S_o_ = S_A_ * S_S_.

### Sequence similarity and interaction profile

To investigate the relationship between the similarity of effector sequences and the similarity of their interaction profiles we calculated the pairwise Jaccard index, which measures the overlap between two effectors’ interaction profiles. We calculated the Jaccard index of all possible effector pairs within a homology cluster. This index represents the ratio of number of human proteins targeted by both effectors to the total number of human proteins targeted by either of them. For our analysis, we only considered effector pairs where the total number of human proteins that are targeted by either effector was at least 3. We implemented the calculations described here as commands in R version 4.2.1.

### Interface predictions

We used as input a representative set of effectors identified in isolated strains (2300 sequences clustered at 90% sequence identity) and all effectors identified in MAGs (186). We ran *mimic*INT as described in^32^ and available at [https://github.com/TAGC-NetworkBiology/mimicINT]. Briefly, mimicINT performs domain searches in effector sequences with InterProScan^90^ using the domain signatures from the InterPro database^91^ retaining matches with an E-value below 10^-5^. For host-like motif detection, mimicINT uses the SLiMProb tool from the SLiMSuite software package^92^ by exploiting the motif definitions available in the ELM database^93^. Motifs are detected in disordered regions as defined by the IUPred algorithm^94^ using both short and long models (motif disorder propensity = 0.2, minimum size of the disordered region = 5). The interface inference step relies on the 3did database^95^ (ii) the ELM database^93^. The workflow checks whether any of the effector proteins contains at least one domain or motif for which an interaction template is available. In this case, it infers the interaction between the given protein and all the host proteins containing the cognate domain (i.e., the interacting domain in the template). To control for false positive inference using motif-domain templates, mimicINT provides two scoring strategies. First, considering binding specificity of domains belonging to the same group (as PDZ or SH3)^96^ an HMM-based domain score^97^ is computed used to rank or filter the inferred interactions. Second, given the degenerate nature of motifs^98^, mimicINT, using Monte-Carlo simulations, assesses the probability of a given SLiM to occur by chance in query sequences and, thus, can be used to filter false positives^99^. This statistical approach randomly shuffles the disordered regions of the input sequences to generate a large set of N randomized proteins.

Here, we first grouped effectors sequences by strain and effectors from MAGs were assigned to the closest strain. In the first experiment, disordered regions were shuffled 100,000 times using as background the effector sequences from the same strain (within-strain shuffling). In the second, regions were shuffled 100,000 times using as disorder background the full set of effector sequences (inter-strain shuffling). Subsequently, the occurrences of each detected motif in each effector sequence were compared to the occurrences observed in the corresponding set of shuffled sequences. We considered as significant all the motif occurrences having an empirical *P* value lower than 0.1. To evaluate whether the number of interface-resolved interactions inferred by mimicINT is significantly different from chance, we generated 10,000 random networks by sampling human proteins from the interaction search space in a degree-controlled manner. We then counted how many randomly generated networks mimicINT inferred a higher number of interfaces than for the one observed in the main screen network. Results and statistical details are in Extended Data Table 3.

### Holdup assay

Domain production: 54 human PDZ domains and the 11 tandem constructs were recombinantly expressed as His_6_-MBP-PDZ constructs in *E. coli* BL21(DE3) pLysS in NZY auto-induction LB medium (nzytech, MB17901)^100^. PDZ domains were purified by Ni^2+^-affinity with a 96-tip automated liquid-handling system (Tecan Freedom Evoware) using 800 µl of Ni^2+^ Beads (Chelating Sepharose Fast Flow immobilized metal affinity chromatography, Cytiva) for each target. The domains were eluted in 2.5 ml of elution buffer: 250 mM imidazole, 300 mM NaCl, 50 mM Tris, pH 8.0 buffer, and then desalted using PD10 columns (GE healthcare, 17085101) into 3.5 ml of 50 mM Tris, pH 8.0, 300 mM NaCl, 10 mM Imidazole buffer. Concentration of desalted His_6_-MBP-PDZ was determined using absorption at 280 nm on a PHERAstar FSX plate reader (BMG LABTECH). Stock solutions were diluted to 4 µM and frozen at −20°C. To assess purity and confirm the concentrations, proteins were further analyzed by SDS-PAGE (LabChip™ GXII, Perkin Elmer). Peptides: 10-mers corresponding to the C-terminal sequences of effectors were ordered as synthetic biotinylated peptides from GenicBio Limited (Shanghai, China); the N-terminal biotin was attached via a 6-aminohexanoic acid linker, which we showed does not alter the peptide’s binding or structural properties^34^. Purity was assessed by HPLC and mass spectrometry; all peptides were >95% pure. Depending on the amino acid composition and charge peptides were solubilized in dH_2_O, 1.4% ammonia or 5% acetic acid, aliquoted at 10 mM concentration and stored at −20°C.

For the hold-up assay we followed published procedures^34,35^. Briefly, 2.5 µl of Streptavidin resin (Cytiva, 17511301) were incubated for 15 min with 20 µl of a 42 µM biotinylated peptide solution, in each well of a 384-well MultiScreenHTS™ filter plate (Millipore, MZHVN0W10). The resin was washed with 10 resin volumes (resvol) of hold-up buffer (50 mM Tris HCl, 300 mM NaCl, 10 mM imidazole, 5 mM DTT), and depleted by incubation for 15 min with 5 resvol of a 1 mM biotin solution, and three washes with 10 resvol of hold-up buffer. A single PDZ domain was then added to each well, incubated for 15 min with the peptide bound to the resin and the unbound PDZ was recovered by centrifugation into 384-well black assay plates for fluorescence readout. The concentration is quantified by intrinsic Trp fluorescence, fluorescein/mCherry was used for peak normalization. Binding affinities and equilibrium dissociation constants (k_D_) were calculated as in^34^, using the mean PBM concentration for k_D_ calculations. Raw values and statistical analysis are in Extended Data Table 3.

### Fluorescent polarization

All FITC labelled peptides were synthesized as 10-mers by Biomatik, Canada, as acetate salts of >98% purity. The FP experiments were performed with the His_6_-MBP-PDZ proteins in 50 mM Tris, 300 mM NaCl, 1 mM DTT, pH 7.5 buffer in 384-well plates (Corning 3544). For direct binding the His_6_-MBP fused PDZ domains were two-fold serially diluted with 12 dilutions, and a final volume of 10 µl. These were then incubated with 50 nM of the FITC labelled viral peptides and the plates were then read out after 1 h in FlexStation 3 (Molecular Devices) at 23°C, using 485 nm excitation and 520 nm emission. For competition experiments, the PDZ domain and FITC peptide were kept constant at 6 µM and 50 mM, respectively. The bacterial effectors peptides in 1% ammonia buffer were added to the PDZ in a four-fold dilution, (5 concentrations: 0 to 31.25 µM) and incubated at room temperature for 2 h. The FITC peptides were then added and further incubated for 1 h at RT. The plates were then read as above. Statistical analysis was performed using the Kruskal-Wallis test with Dunn’s test followed by an FDR-correction. Raw values and statistical analysis are in Extended Data Table 3.

### Effector convergence

To estimate the significance of effector convergence, we performed a permutation test by randomly sampling ‘target’ nodes (n = 979) from Y2H identifiable proteins from the human reference interactome map, HuRI^86^, as the sampling space (n *=* 8,274). We used sampling with replacement to allow repeatedly picking a protein. In each iteration, the number of distinctly targeted proteins was counted. The resulting distribution from 10,000 random permutations was used to calculate the z-score of the experimentally observed targets (n = 349). The *P* value is the area under the curve for the standard normal distribution up to a given z-score. We calculated the *P* value as implemented in the “pnorm()” R function using the z-score as input. To account for the two-tailed test, the *P* value was multiplied by 2. To avoid artifacts due to differential sampling we only considered interactions in the HuMMI_MAIN,_ excluding those human proteins targeted by effectors of the unknown strains and targets outside HuRI. The rationale for the latter is that a substantial proportion of proteins that are not in HuRI may not be suitable for Y2H analysis. Thus, restricting the analysis to the HuRI subset increases the stringency.

To estimate the significance of the convergence of effectors from different strains (interspecies convergence), we used a conditional permutation test that preserves the strain contribution. For each iteration, we generated 18 samples, where for each sample, we randomly picked the number of proteins equivalent to the observed targets of each strain (Extended Data Table 3). From the full list of random picks that are assigned to all strains, the frequency of selecting a protein was recorded. This frequency is the convergence value which indicates the number of targeting strains. Using the convergence value distribution obtained from 10,000 iterations, we identified the statistically significant number of strains sharing a target. The observed convergence value ranges from 2 to 15 strains. We calculated the z-scores using the convergence value distribution obtained from the conditional permutation test and the associated *P* values as implemented in the “*pnorm()*” R function. The significant convergence value (*P* value < 0.004) starts at 4 strains. We considered any target that is in common between at least 4 strains to be subject to interspecies convergence.

### Function enrichment analysis

We used the “*gost()*” function from the gprofiler2 version 0.2.1 R package^101^ to identify enriched functions in effector targets. This function implements a hypergeometric test to estimate the significance of the abundance of genes considering the frequency of the genes in the function annotation databases. The main input argument for this function is the gene list (“*query*”). The function allows the user to optionally set input arguments, including the background (“*custom_bg*”), evidence codes (“*evcodes*”), annotation databases (“sources”), methods for correcting the hypergeometric test *P* values (“*correction_method*”), and other arguments that were set to their default options. We used the target official symbol identifiers as the “query” argument. The list of HuRI proteins was the “*custom_bg*” argument. The annotations inferred from electronic annotations were excluded by setting the “*exclude_iea*” argument to “*TRUE*”. The hypergeometric test *P* values were corrected using Benjamin-Hochberg method by setting the “correction_method” argument to ”*fdr*”. The argument (“*sources*”) was set to a vector (“*GO:BP*”, “*KEGG*”,”*REAC*”), which encodes the search space across three function annotation databases: gene ontology biological process terms (“*GO:BP*”)^102^, Kyoto encyclopedia of genes and genomes (“*KEGG*”) pathways^103^, and Reactome pathway database (“*REAC*”)^104^. After plugging in these inputs into the “*gost()*” function, the output is a named list where “result” is a data frame that tabulates the enrichment analysis results. We calculated the odds ratio and the fold enrichment to estimate the effect size of each tested function. The odds ratio was calculated for each function as the odds in the target set divided by the odds in the HuRI set. The odds in the target set are the number of function-annotated target proteins divided by that of the function-unannotated target proteins. Similarly, the odds in the HuRI set are the number of function-annotated HuRI proteins divided by that of function-unannotated HuRI proteins. The fold enrichment was calculated for each function by comparing the number of function-annotated target proteins to that of the expected. The expected value represents the number of function-annotated target proteins that is expected randomly based on the HuRI background. It is the product of the total number of targets (n = 349) by the rarity. The rarity is the number of function annotated HuRI proteins divided by the sum of annotated HuRI proteins. The total HuRI proteins annotated for GO:BP, KEGG, and REAC, are 6988, 3250, and 4592, respectively. Statistical details are in Extended Data Table 5.

### Metabolic subsystem analysis

Several metabolism-related functions were significantly enriched in target proteins; therefore, we tested the abundance of targeted enzymes in metabolic subsystems using the human genome-scale metabolic model Recon3D^46^. To focus on metabolic enzymes as opposed to signaling enzymes, we excluded ligases and kinases from Recon3D analyses. We performed the hypergeometric test using the R function “*phyper()*” for each subsystem annotated in Recon3D (n = 95). The inputs to this function are: the number of subsystem-annotated targeted enzymes, the number of subsystem-annotated Recon3D enzymes, the number of subsystem-unannotated Recon3D enzymes, and the number of targeted enzymes (n = 16). The nominal *P* values were corrected using Benjamin-Hochberg. We calculated the odds ratio and the fold enrichment using the same calculations described above for functional enrichments.

### Random walk-based determination of commensal effector network neighborhoods

We have implemented a network propagation protocol based on a Random Walk with Restart (RWR) algorithm RWR-MH^105^ to explore the network vicinity of the commensal effectors in HuRI^54^, which contains 338 target proteins (HuMMI_MAIN_ screen) of 243 commensal effectors. We used the human effector targets as seeds for the random walk and set the restart probability to the default value of 0.7. In this way, we obtained a ranked list of proteins in the network: the ones with the higher scores are more proximal to the seeds than those with lower scores. To assign statistical significance to the computed RWR scores, we implemented a normalization strategy based on degree-preserving network randomizations^106^. We thus generated 1,000 random networks from HuRI and ran the RWR algorithm to compute 1,000 scores for each network protein. We then computed an empirical *P* value for each protein in the network keeping as neighbor proteins only those with an empirical *P* value < 0.01.

### Disease enrichment analysis

We tested the association of all target proteins, or those subject to convergence, with human diseases by performing a two-sided Fisher’s exact test. We used the disease-causal genes identified by the Open Targets genetic portal, which prioritizes genes at GWAS loci based on variant-to-gene distance, molecular QTL colocalization, chromatin interaction, and variant pathogenicity^107^. This machine-learning approach assigns a locus to gene (l2g) score to identify the most likely causal gene for the genetic variation signal of any marker SNP. We considered a score of 0.5 or more as a threshold, as recommended by the authors^108^. The Fisher’s exact test was performed using the function “*fisher.test()*” from “*stats*” R package version 4.2.2 with its default inputs whenever applicable. The input to this function is a 2 x 2 contingency table, where columns represent the query set and the background set, and rows denote the absence or presence of causal genes in the respective set. HuRI proteins were used as the background set, and the query set was either the target proteins or those subject to convergence. The calculated nominal *P* values from this function were then corrected using the Benjamin-Hochberg method as implemented in the “*p.adjust()*” function. The odds ratio and fold enrichment values were calculated as described in the functional enrichment section. Statistical details are in Extended Data Table 5.

### Association with human traits and phenotype in network neighborhoods

For each set of significant neighborhood-proteins we tested for enrichment of Open Targets causal genes for human traits that had been investigated by 3 or more studies and for which the Open Targets initiative identified 3 or more causal genes (l2g ≥ 0.5). We used a two-sided Fisher’s exact test to assess whether a given strain neighborhood is enriched in protein associated with a human trait or phenotype followed by Benjamini-Hochberg multiple testing correction. This yielded no significant association (FDR < 0.05). We therefore focused on 400 associations with a nominal *P* value < 0.01 and an OR > 3. Some disease categorizations were adjusted to better reflect etiology. Thus, Sjogren syndrome, eczema and psoriasis were considered an ‘immunological’ rather than eye or skin traits, and osteoarthritis was labeled as a disease of “musculoskeletal or connective tissue” rather than metabolic. For Fig. 4d some closely related traits were merged, i.e., three asthma terms and three psoriasis terms. Statistical details are in Extended Data Table 5.

### NF-κB activation assay

HEK 293 (RRID: CVCL_0045, DSMZ) were maintained in DMEM with 10% FBS and 100 U/mL penicillin and 100 U/mL streptomycin at 37°C and 5% CO2. IKKβ (in pRK5 with a Flag-tag) served as positive control whereas A20 (in pEF4 with a Flag-tag) as the negative control. In a 60 mm cell culture dish 1 x 10^6^ cells were seeded in 3 ml Medium. After 24 h cells were transfected using 10 ng NF-κB reporter plasmid (6 × NF-κB firefly luciferase pGL2), 50 ng pTK reporter (renilla luciferase) and 2 µg bacterial ORF in pMH-FLAG-HA. The DNA was added to 200 µl 250 mM CaCl_2_ solution (Carl Roth cat. no. 5239.1), vortexed and added dropwise to 200 μl 2 × HBS (50 mM HEPES (pH 7.0) (Carl Roth cat. no. 9105.4), 280 mM NaCl (Carl Roth cat. no. 3957.2), 1.5 mM Na_2_HPO_4_ × 2 H_2_O (Carl Roth cat. no. 4984.1, pH 6.93) which was vortexed. After 15 min incubation, the mixture was added dropwise to the cells. Medium was changed after 6 h incubation. To assess NF-κB inhibition, cells were treated for 4 h with 20 ng/ml TNF (Sigma-Aldrich cat. no. SRP3177) 24 h after transfection. Samples were washed, lysed, centrifuged and the supernatant was measured using the dual luciferase reporter kit (Promega, E1980) with a luminometer (Berthold Centro LB960 microplate reader, Software: MikroWin 2010). NF-κB induction was determined as Firefly luminescence to Renilla luminescence. *P* values were calculated using the Kruskal-Wallis test with Dunn’s correction followed by an FDR-correction. Raw values and statistical analysis are in Extended Data Table 6.

Protein expression levels were checked by Western Blots. Proteins were separated by SDS-PAGE and transferred on polyvinylidene fluoride membranes, and after transfer blocked with 5% milk in 1 × PBS + 0.1% Tween-20 (PBST) for 1 h at room temperature. Primary antibodies were added in 2.5% BSA in PBS-T buffer at 4°C overnight. After 3 x 15min washes with PBS-T anti-mouse secondary antibody was added at a 1:10,000 dilution for 1 h at RT (Jackson ImmunoResearch Labs cat. no. 715-035-150, RRID:AB_2340770). Primary antibodies: anti-Actin beta (SCBT cat. no. sc-47778, RRID:AB_626632) at a 1:10,000 dilution, anti-FLAG M2 (Sigma Aldrich cat. no. F3165, RRID:AB_259529) at a 1:500 dilution and anti-HA (Sigma-Aldrich cat. no. 11583816001, RRID:AB_514505) at a 1:1,000 dilution. For detection the LumiGlo reagent (CST cat. no. 7003S) and a chemiluminescence film (Sigma-Aldrich cat. no. GE28-9068-36) were used.

### ICAM1 assay

Caco-2 cells were maintained in DMEM Glutamax medium (Gibco) supplemented with 10% FBS, 1% Pen/Strep at 37°C in a humidified 5% CO2 incubator. Medium was refreshed twice a week. Caco-2 cells were plated in both 24- and 96-well plates 24 h before transfection. Six hours prior to transfection, culture medium was replaced with supplement-free DMEM. Co-transfections were performed using 40,000 MW linear polyethylenimine (PEI MAX®) (Polysciences, Warrington, USA) at a ratio of 1:5 pDNA:PEI. Equimolar ratios of the eGFP-plasmid and effector-plasmid were used to ensure equimolar representation of relevant ORFs. In total, 250 ng and 1 µg pDNA was added per well of the 96- and 24-well plates, respectively. pDNA-PEI complexes were formed by incubating pDNA and PEI at RT for 15 minutes, followed by the addition of supplement-free DMEM and another incubation of 15 minutes at RT. Cells were then exposed to the transfection mixture for 16 h, washed, and rested for 6 h in complete DMEM. Subsequently, cells were stimulated using an activation mix containing 200 ng/ml PMA (P8139-1MG, Sigma-Aldrich), 100 ng/ml LPS (L6529-1MG, Sigma-Aldrich), and 100 ng/ml TNF (130-094-014, Miltenyi Biotec). In 24-well plates, cells were stimulated for 24 h and detached from the plate using ice-cold PBS. In the 96-well plate, cells were stimulated for 48 h, treated with BD GolgiStop™ (554724, BD Biosciences) in the final 6 h of stimulation, and detached using trypsin/EDTA. Cells were washed twice and ICAM1 was stained using an anti-ICAM1 PE (#MHCD5404-4, Invitrogen) antibody. The mean fluorescent intensity of the GFP+ cell population was measured on a FACSFortessa™ flow cytometer (BD) and the data was analyzed using FlowJo V10.8.1 (BD). After positive tests for normal data distribution, significance was assessed using a one-way ANOVA with Dunnett’s multiple comparisons test. Raw values and statistical analysis are in Extended Data Table 6.

### Cytokine assays

Caco-2 cells were plated in 100 mm cell culture dishes three days prior to transfection. The transfection protocol was identical to that described above, however, a total of 20 µg pDNA was used per dish. Upon overnight transfection, cells were detached using Trypsin/EDTA and resuspended in cell sorting buffer (PBS + 2% FBS + 2mM EDTA). GFP+ cells were sorted into ice-cold FBS using a BD FACSAria III cell sorter (BD) and transferred to a 96-well plate at 30,000 cells per well. Upon a 24 h rest-period, cells were activated for 48 h using the activation mix described above. During cell stimulation, cell proliferation was monitored through longitudinal imaging of cell confluency in the Incucyte S3 Live cell analysis system (Essen BioScience). Cytokine levels were determined using the human inflammation panel 1 LEGENDplex™ kit (Biolegend) following the manufacturer’s instructions. Cell culture supernatant of the above samples was used to analyze IL1beta. To this end, IL1beta ELISAs were performed using the ELISA MAX™ Deluxe Set Human IL1beta kit (437015, Biolegend) following the protocol provided by the manufacturer. Statistical significance was evaluated using Kruskal-Wallis test with uncorrected Dunn’s test. Raw values and statistical analysis are in Extended Data Table 6.

### Protein ecology

Metagenomic assemblies from the Inflammatory Bowel Disease Multi’omics DataBases (IBDMBD)^64^ and from the skin metagenome^109^ were downloaded, and each samples protein repertoire predicted using Prodigal (options; -p meta)^110^. Effector proteins were compared to the metagenomic protein repertoires using DIAMOND (options; >90% query length, >80% identity). For analyses in Fig. 5, samples were grouped into patients with UC (n = 304), CD (n = 508), and controls without IBD (n = 334). The annotations were then converted into binarised vectors of presence and absence of each effector across the sample and the Fischer exact test, implemented within scipy python module, was used to determine if the prevalence of each effector occurring within CD or UC patient metagenomes compared to controls. Significance was then corrected using the Benjamini-Hochberg method. The significance of differences in prevalence distributions between healthy and either patient cohort were estimated by Wilcoxon rank-sum test, implemented in the “*wilcox.test()*” R function. Statistical details in Extended Data Table 6.

### Statistics and reproducibility

Data were subjected to statistical analysis and plotted to Microsoft Excel 2010 or python or R scripts. For comparison of normally distributed values we used one-way ANOVA, for assessment of overlap for comparison of values not passing the normality tests we used Kruskal-Wallis test with Dunn’s corrected as appropriate and indicated in the figure legends and methods. Enrichments were calculated using Fisher’s exact test with Bonferroni FDR correction. All statistical evaluations were done as two-sided tests. Generally, a corrected *P* value < 0.05 was considered significant. GO, KEGG, and Reactome functional enrichments were calculated using profiler with the respectively indicated background gene sets. For the disease target enrichments and neighborhood associations no associations were significant after multiple hypothesis correction, which is why nominally significant associations calculated by Fisher’s exact tests were used for Fig. 4c,d. All raw values, n, and statistical details are presented in supplementary tables as indicated in the Figure legends and methods sections.

## AUTHOR CONTRIBUTIONS

Project conception: PFB

T3SS and effector analyses: PH, TH, SA, CB, AZ, TR, PFB

ORF cloning: VY, MR, MA, AS, PFB

Interactome mapping and validation: VY, SR, BW, AS, PFB

Interaction curation: VY, MA, MB, AZ, CF, PFB

Data analyses: BD, VY, DS, CWL, MB, SAC, PS, CB, AZ, PFB

Interface identification and validation: SAC, AZ, JFM, SBM, JCT, RV

Effector ecology: TH, TC

Cell-based assays: VY, NvdH, FO, PFB, DK, MB

Visualization: VY, BD, JFM, AZ, PFB

Funding acquisition: PFB, CF, AZ, CB, TR, DK

Manuscript writing and editing: PFB, VY, BD, BW, TH, CF, AZ

## ACKNOWLEDGEMENTS

Plasmids and strains for hsPRS/RRS_v2 were kindly provided by Marc Vidal, David E. Hill, and Mike Calderwood, CCSB, Dana-Farber Cancer Institute, Boston, MA. The computational results presented have been achieved in part using the Vienna Scientific Cluster (VSC). Centre de Calcul Intensif d’Aix-Marseille is acknowledged for granting access to its high-performance computing resources.

## REPORTING SUMMARY

Further information on research design is available in the Nature Research Reporting Summary linked to this paper.

## DATA AVAILABILITY

All sequence, interaction, and functional data generated in this study are available as supplementary information. The effectors identified and cloned for interactome mapping are presented in Extended Data Table 1. All protein-protein interaction data acquired in this study can be found in Extended Data Table 2 and Extended Data Table 3. The data for functional validation assays can be found in Extended Data Table 6. The protein interactions from this publication have been submitted to the IMEx (http://www.imexconsortium.org) consortium through IntAct^111^ and assigned the identifier IM-29849. New effector sequences have been submitted to GenBank: BankIt2727690: OR372873 - OR373035 and OR509516 - OR509528.

## CODE AVAILABILITY

All source code related to this paper is available as a zip file.

## EXTENDED DATA TABLES

Extended Data Table 1: T3SS in strains of the commensal human microbiome

Extended Data Table 2: Effector identification and cloning

Extended Data Table 3: Effector host interaction map

Extended Data Table 4: Interface identification and validation

Extended Data Table 5: Functional and disease enrichment

Extended Data Table 6: Functional assay data and IBD prevalence

**Extended Data Fig. 1.**
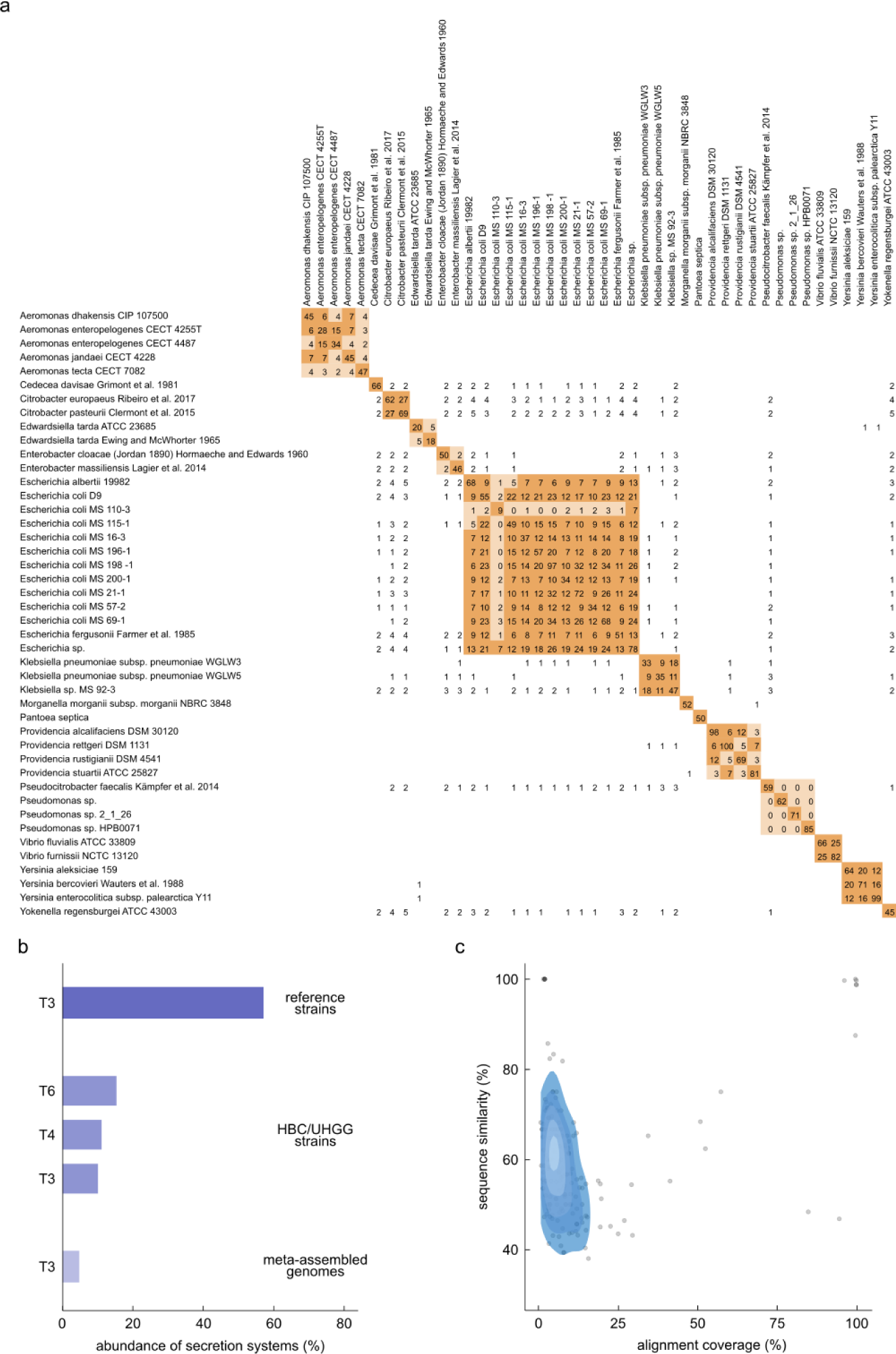
| T3SS in strains of the commensal gut microbiome. **a**, Effector-complement comparison of the 44 T3SS+ Pseudomonadota reference strains. Numbers indicate the count of shared effectors at >90% mutual sequence similarity across 90% common sequence length among the indicated strains. **b**, Abundance of secretion systems in Pseudomonadota genomes among the 77 reference strains of human intestinal and stool samples, in a collection of 4,475 strains isolated from normal human guts (HBC/UHGG strains) and in meta-assembled genomes (MAG) of normal human guts. **c**, Similarity of identified 186 candidate effectors from the 770 T3SS+ MAGs with 1,195 effectors from pathogenic microbes across the range of alignment coverages. Full data for all panels in Extended Data Table 1.

**Extended Data Fig. 2.**
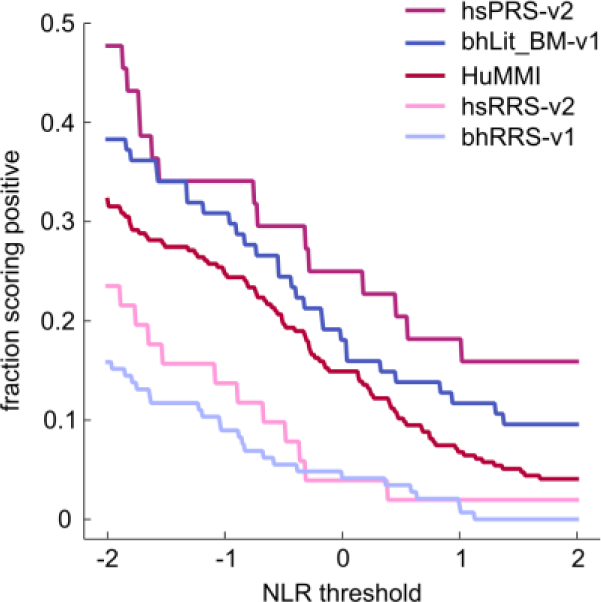
| Detection rates of protein pairs in different sets across varying thresholds in yN2H. Fractions scoring positive of the HuMMI dataset and benchmarking datasets (hsPRS-v2, bhLit_BM-v1, hsRRS-v2, bhRRS-v1) depending on the threshold of the normalized luminescence ratio (NLR). Full data in Extended Data Table 3.

**Extended Data Fig. 3.**
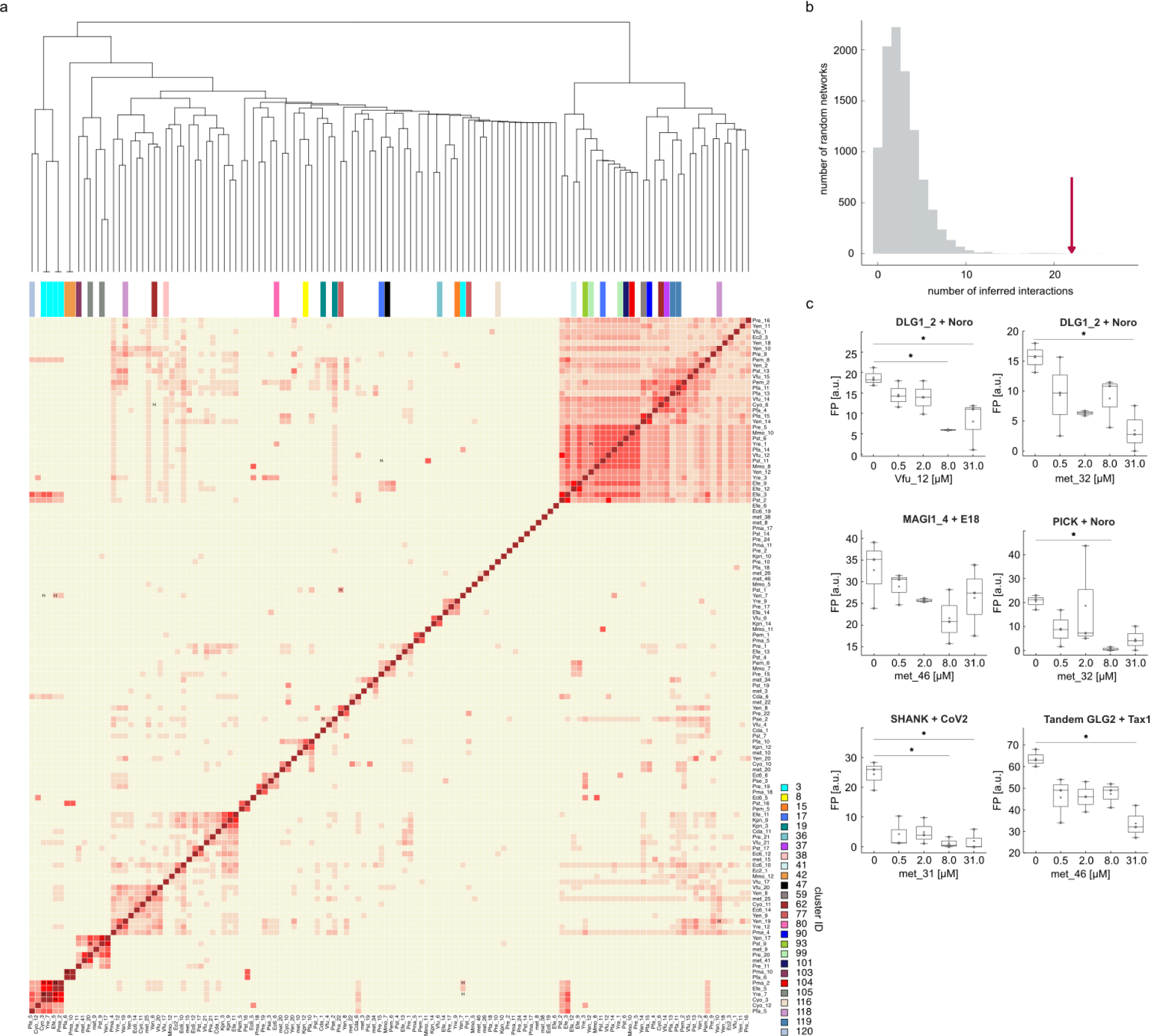
| Interaction specificity and interaction motifs. **a**, Jaccard-interaction similarity of all interacting effector-pairs with at least 3 shared human interactors. Color-intensity correlates with Jaccard-index. Effector pairs marked with “H” share the same homology cluster. Clusters are color-coded according to legend. **b**, Count of motif-domain pairs matching at least two stringency criteria identified in HuMMI_MAIN_ (arrow) compared to n = 10,000 randomized control networks (empirical *P* = 0.0003). **c**, Competition of the interaction between human PDZ domain and viral PBM peptide by indicated C-terminal effector peptides. * *P* < 0.05 (Kruskal Wallis with Dunn’s correction, n = 3). Boxes indicate IQR, black line represents mean, whiskers indicate highest and lowest data point within 1.5 IQR. Precise *P* values and n for each test are shown in

**Extended Data Fig. 4.**
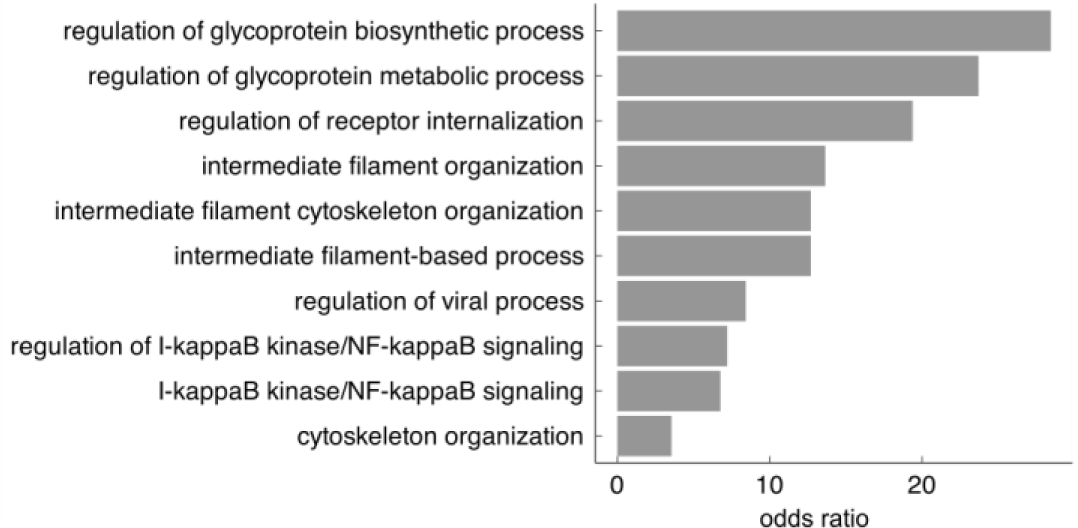
| GO enrichment for convergence proteins. OR for functional annotations enriched among effector-targeted human proteins that are subject of convergence (FDR < 0.05, Fisher’s exact test with Bonferroni FDR correction). Full data and precise FDR and OR values in Extended Data Table 5.

**Extended Figure 5.**
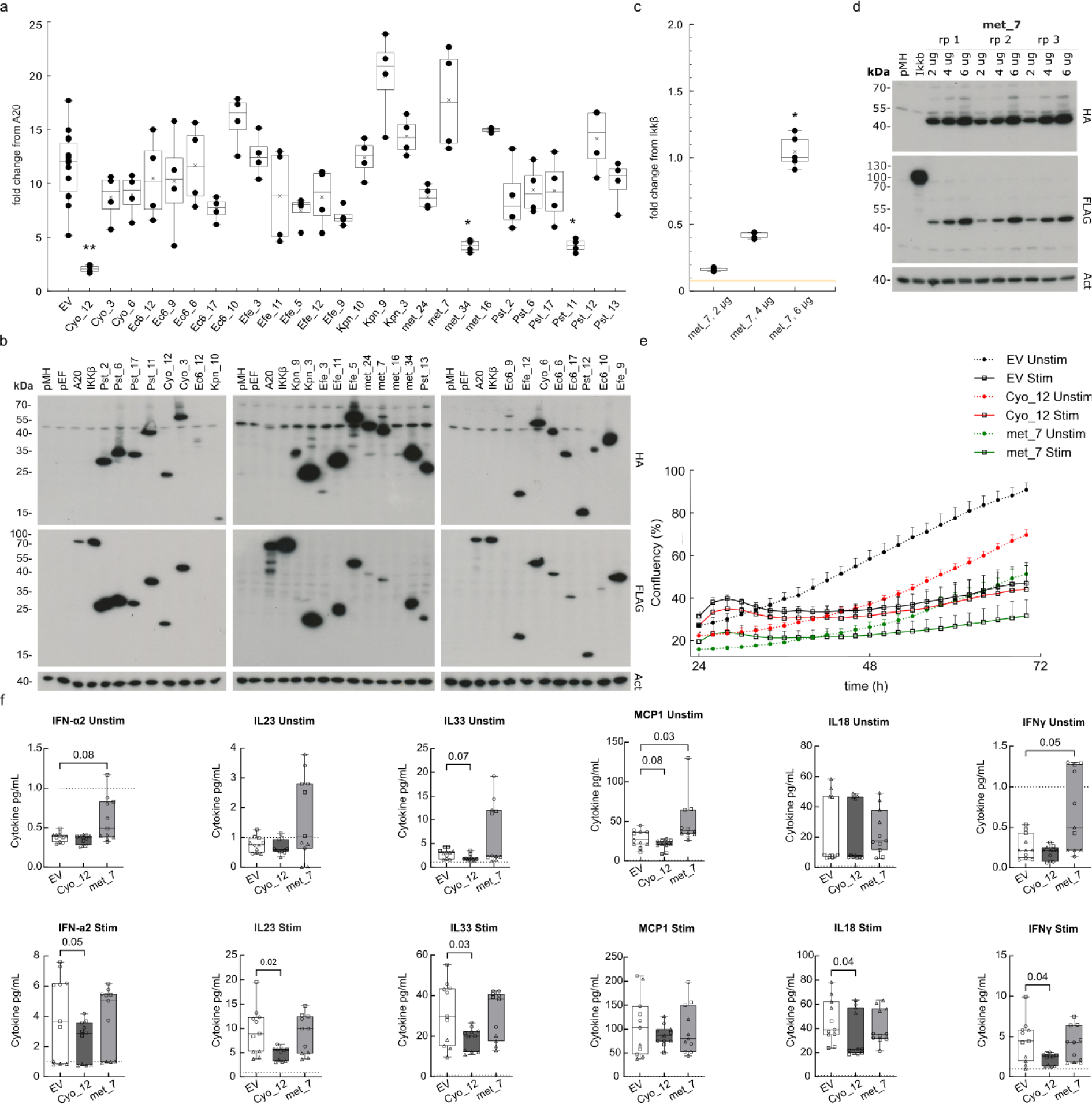
| Effector impact on human cell function. **a.** Relative NF-κB transcriptional reporter activity of HEK293 cells expressing the indicated effectors under TNF-stimulated conditions (Kruskal-Wallis test with Dunn’s correction, * *P <* 0.05, ** *P* = 0.01, n = 4). Boxes represent IQR, with the bold black line representing the mean; whiskers indicate highest and lowest data point within 1.5 IQR. **b**, Representative anti-Hemagglutinin (HA) and anti-Flag (FLAG) western blots showing expression of transfected effector proteins relative to actin control (ACT). Empty pMH-Flag-HA (pMH), empty pEF4 (pEF). **c**. Titration of met_7 shows a concentration dependent specific increase of NF-κB reporter activity. Yellow line represents the empty vector value. (Kruskal-Wallis test with Dunn’s correction, * *P <* 0.05, error bars: standard deviation of the mean, n = 5). Boxes represent IQR, with the bold black line representing the mean; whiskers indicate highest and lowest data point within 1.5 IQR. **d**, Representative anti-Hemagglutinin (HA) and anti-Flag (FLAG) western blots for experiment in c showing expression of transfected effector proteins relative to actin control (ACT). **e**, Representative proliferation curves of Caco-2 cells transfected with empty vector (EV), Cyo_12 or met_7 in basal conditions (unstim) or following pro-inflammatory stimulation (stim) over 72 h after sorting. **f**, Concentration of cytokines secreted by Caco-2 cells transfected with the indicated effectors in basal conditions (Unstim) or following pro-inflammatory stimulation (Stim). EV indicates empty vector mock control. Indicated *P* values calculated by Kruskal-Wallis test with Dunn’s multiple hypothesis correction (n = 11). Boxes represent IQR, with the bold black line representing the mean; whiskers indicate highest and lowest data point. Raw measurements, n, and precise *P* values for all panels in Extended Data Table 6.

